# The circadian hippocampus and its reprogramming in epilepsy: impact for chronotherapeutics

**DOI:** 10.1101/199372

**Authors:** K. J. Debski, N. Ceglia, A. Ghestem, A. I. Ivanov, G. E. Brancati, S. Bröer, A. M. Bot, J. A. Müller, S. Schoch, A. Becker, W. Löscher, M. Guye, P. Sassone-Corsi, K. Lukasiuk, P. Baldi, C. Bernard

## Abstract

Gene and protein expression displays circadian oscillations in numerous body organs. These oscillations can be disrupted in diseases, thus contributing to the disease pathology. Whether the molecular architecture of cortical brain regions oscillates daily and whether these oscillations are modified in brain disorders is less understood. We identified 1200 daily oscillating transcripts in the hippocampus of control mice. More transcripts (1600) were oscillating in experimental epilepsy, with only one fourth oscillating in both conditions. Proteomics confirmed these results. Metabolic activity and targets of antiepileptic drugs displayed different circadian regulation in control and epilepsy. Hence, the hippocampus, and perhaps other cortical regions, shows a daily remapping of its molecular landscape, which would enable different functioning modes during the night/day cycle. The impact of this remapping in brain pathologies needs to be taken into account not only to study their mechanisms, but also to design drug treatments and time their delivery.

---

Circadian rhythms regulate numerous brain functions (e.g. wakefulness, feeding, etc.) with 24-hour oscillations in most species. The core time keeping machinery is the suprachiasmatic nucleus (SCN). Its molecular architecture undergoes a daily remapping driven by transcriptional and translational feedback loops, thus allowing switching between different functional states ^1, 2^. Several studies suggest the existence of additional (ancillary) clocks in the brain outside the SCN ^3^. Indeed, clock genes, including Per1, Per2, Bmal1, Cry1 and Cry2, exhibit rhythmic expression in a brain region- and gender-dependent manner ^4, 5^. Since these genes are involved in the regulation of transcription/translation of many other genes, brain regions outside the SCN may also undergo a complex daily remapping of their molecular architecture ^6, 7^. Such circadian regulation of neuronal circuits may enable appropriate coordination between physiology and behavior, since learning and memory processes are regulated in a circadian manner ^8, 9^. In the hippocampus, a region critical for numerous learning and memory processes, synaptic responses ^10^, long term synaptic plasticity ^11, 12^ and the memory-associated MAPK pathway ^13^ display circadian rhythmicity.

The first goal of this study is to provide the community with a resource listing of the genes and proteins undergoing circadian regulation in the mouse hippocampus. Such information is crucial not only for our understanding of hippocampal function, but also for a proper interpretation of already published data and for resolving discrepancies, since experiments performed at different times may occur within different molecular landscapes. The second goal of this study is to provide a similar resource in a mouse model of temporal lobe epilepsy (TLE), the most frequent form of epilepsy in adults. Several reasons motivate this choice. Alterations of circadian rhythms can lead to pathologies ^14^, and conversely, disruption of the circadian clock can have direct deleterious consequences ^15, 16^, e.g. on sleep patterns ^17^. In the liver, the circadian clock is reprogrammed by nutritional challenge, resulting in a different cycling of genes and proteins ^18^. Since clock genes display different cycling behavior in experimental epilepsy ^19^, the circadian regulation of downstream genes may follow different rules in control and TLE conditions. This information is essential for a proper interpretation of studies comparing the levels of expression of genes and proteins, a standard experimental procedure to assess the reorganization of neuronal circuits in TLE (and in neurological disorders in general). It is also essential for chronotherapy (the proper timing of drug delivery) ^20^, as many pharmaceutical molecules target gene products showing circadian rhythmicity ^21^.

## RESULTS

### Circadian regulation of genes and proteins in the mouse hippocampus

Since the distribution of transcripts is heterogeneous along the septo-temporal axis of the hippocampus ^22^, we focused on its ventral part ^23^, which is most epileptogenic in human TLE and experimental models ^24^. Using Affymetrix microarrays and cluster analysis, we identified 1256 transcripts oscillating in the ventral hippocampus of control mice (**Fig. 1a** and **Supplementary Table 1**). Hence, 10% of hippocampal genes are regulated in a circadian manner, as compared to 19% in the SCN ^5^. Data from post mortem human hippocampus suggests that less genes (n=659) are oscillating ^7^. Comparison with the latter study is difficult because investigating circadian rhythms requires a strict control of circadian conditions (e.g. no light exposure during the night), which could not be achieved in the human study.

**Figure 1.**
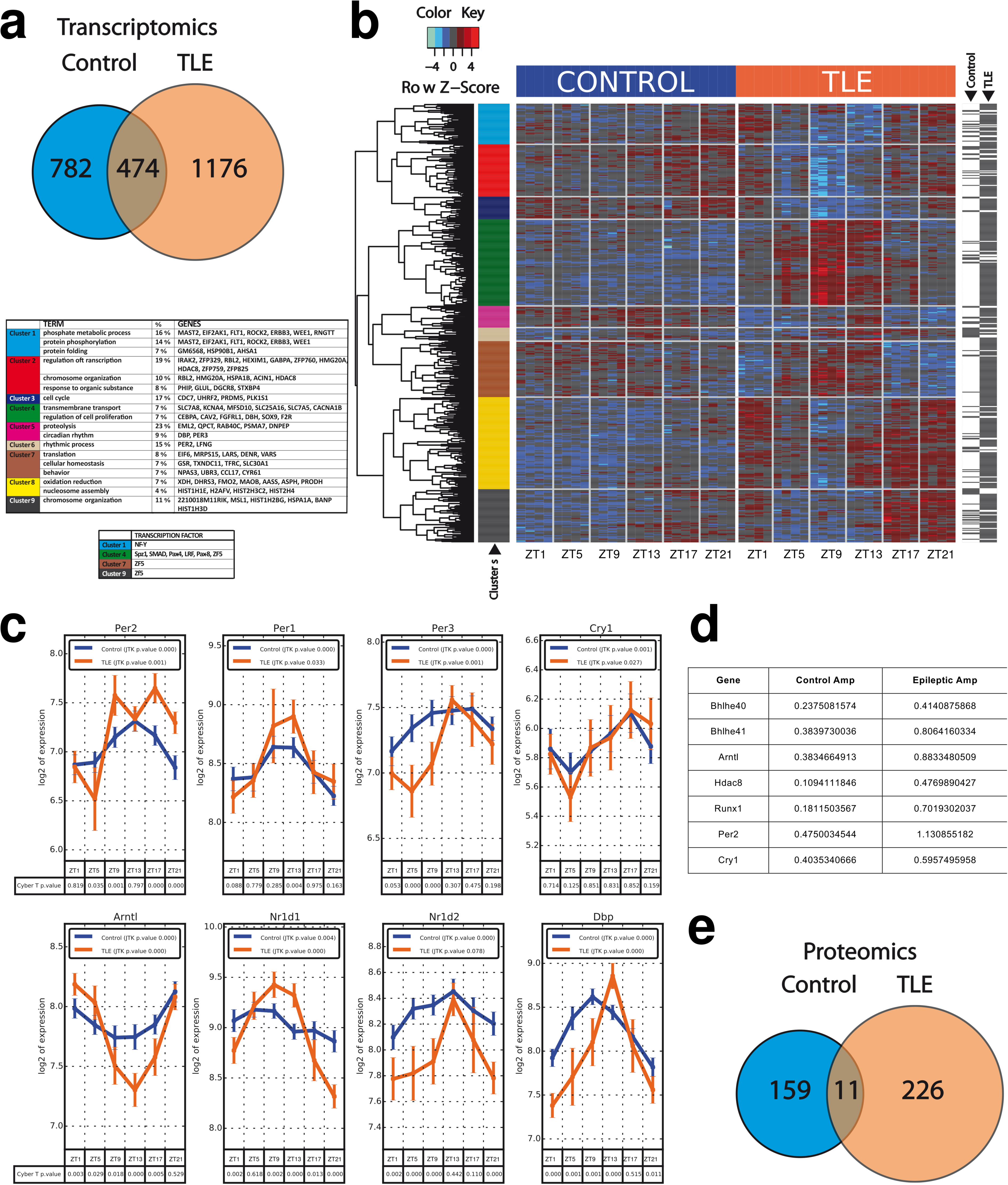
Circadian regulation of genes and proteins in the hippocampus and their reprogramming in epilepsy. (**a**) In control mice (n=4 per Zeitgeber Time – ZT), 1256 transcripts are oscillating. In TLE (n=4 per ZT), more transcripts are oscillating (1650). Only 474 are common to both conditions, showing a remodeling of the landscape of oscillating genes in TLE. (**b**) Right panel. Heatmap of 1105 mRNAs showing circadian regulation. 124 mRNAs were differentially expressed in at least one time point only in control animals, 908 mRNAs were differentially expressed only in TLE and 73 mRNAs were differentially expressed in both groups. Each column per time window represents an individual animal and each row represents an individual mRNA. Colors on the heatmap represent Z-score: higher – red, lower – blue. The hour of tissue collection is indicated below (ZT). The dendrogram obtained from hierarchical clustering is shown on the left side of the heatmap panel. Genes are ordered by clustering complete-linkage method together with Pearson correlation distance measure. Colors in the bar on the left side of the heatmap panel represent clusters obtained by cutting dendrogram at selected level heights to obtain nine groups. Black lines in the bar on the right side of the heatmap panel mark genes showing differences in expression between control or TLE animals based on one-way ANOVA (cut-off FDR < 0.1). Left top panel. Biological functions for each gene cluster, according to Gene Ontology vocabulary using DAVID. Only terms, which are represented by more than 5% of genes in a given cluster, are presented in the Table. Left bottom panel. Transcription factors with binding sites overrepresented in different gene clusters. (**c** and **d**) Oscillation patterns and amplitudes of core clock genes. Note the phase change and general increase in oscillation amplitude in TLE as compared to controls. (**e**) In control mice 170 peptides are oscillating, whilst 337 are oscillating in TLE. Only 11 peptides oscillate in both conditions, indicating a reprogramming of the molecular daily remapping of the hippocampus in TLE.

Using cluster analysis and a false discovery rate < 0.01, we found 197 transcripts differentially expressed in at least one time point, which could be identified based on biological functions (**Fig. 1b**). When compared to subcortical structures (**Table 1**), we found a clear overlap of core clock genes involved in transcriptional and translational feedback loops in the SCN across all investigated brain regions (highlighted in green in **Table 1**). Oscillating core clock genes in the hippocampus included Per1-3, Cry1, Arnt1, Nr1d1-2, and Dbp (**Fig. 1c**, **Table 1**). Transcriptome analysis was confirmed with qPCR (**Supplementary Fig. 1**). Interestingly, a large set of oscillating genes was specific to the hippocampus (**Table 1**), such as Creb1 (**Fig. 2a**), which is involved in both synaptic plasticity and synchronization of circadian rhythms ^25^, demonstrating that circadian rhythmicity of genes is brain region-dependent.

**Figure 2.**
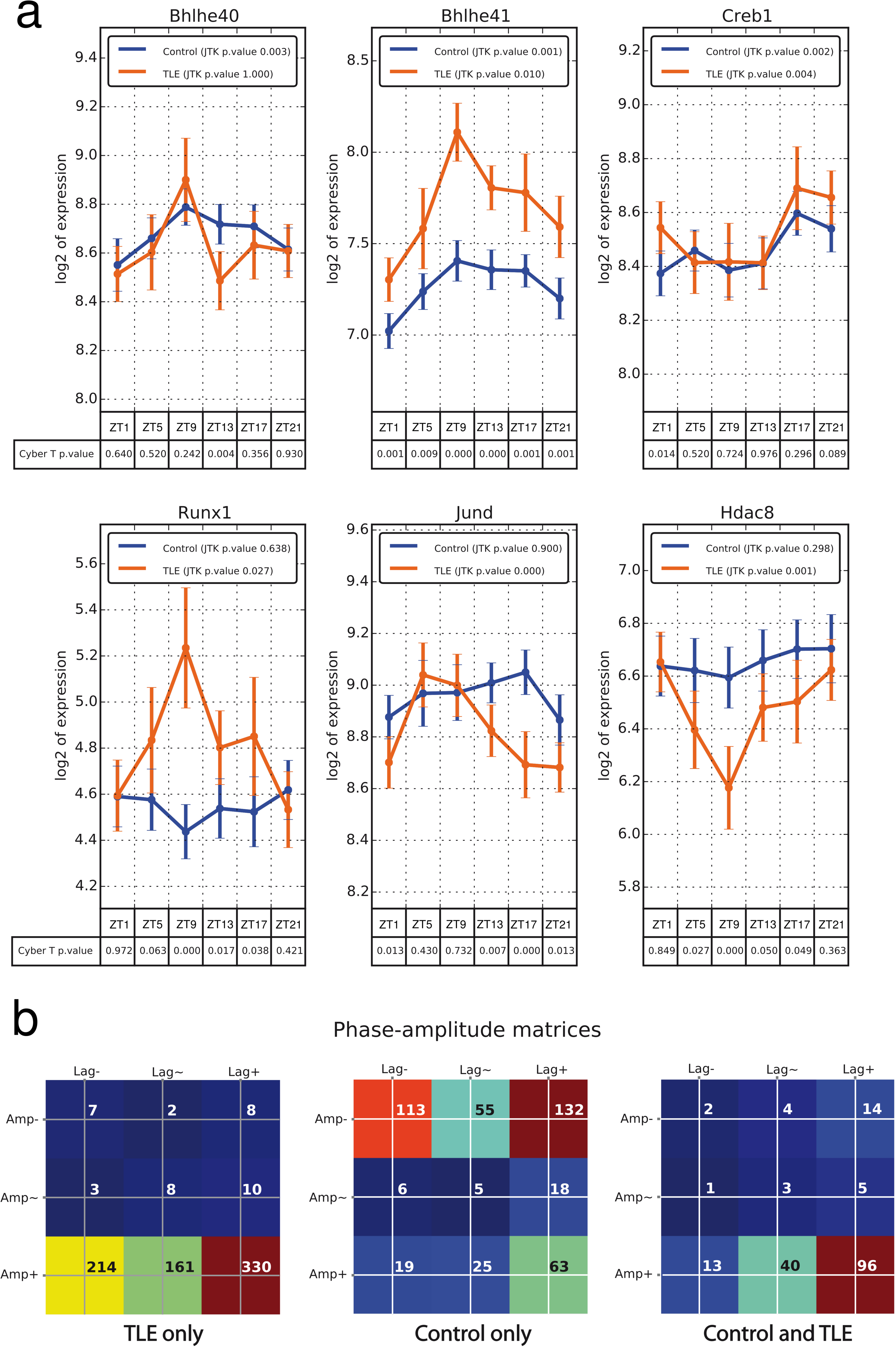
Different oscillatory patterns of gene transcripts in control mice and their alterations in TLE. (**a**) Oscillatory patterns of genes, which may contribute to the reprogramming of the circadian hippocampal remapping in TLE. *Creb1* oscillates in both conditions. Key controllers of circadian rhythms, *Bhlhe40* and *Bhlhe41*, show increased oscillation amplitudes in TLE, which may contribute to the large recruitment of oscillating genes in TLE. *Runx1*, a DNA binding regulator, and *Hdac8*, a chromatin remodeler, gain statistically significant oscillation in TLE. *JunD*, which interacts with the AP-1 transcription factor complex and *Creb1* shows a 180° phase shift in TLE as compared to control. (**b**) Phase-Amplitude matrices. Most of the transcripts show high amplitude oscillations in TLE, whilst they are distributed in a bimodal manner in controls between low and high amplitude. The majority of high amplitude oscillatory transcripts are phase advanced as compared to the rest of the oscillatory transcripts, a property more pronounced in TLE.

**Table 1:**
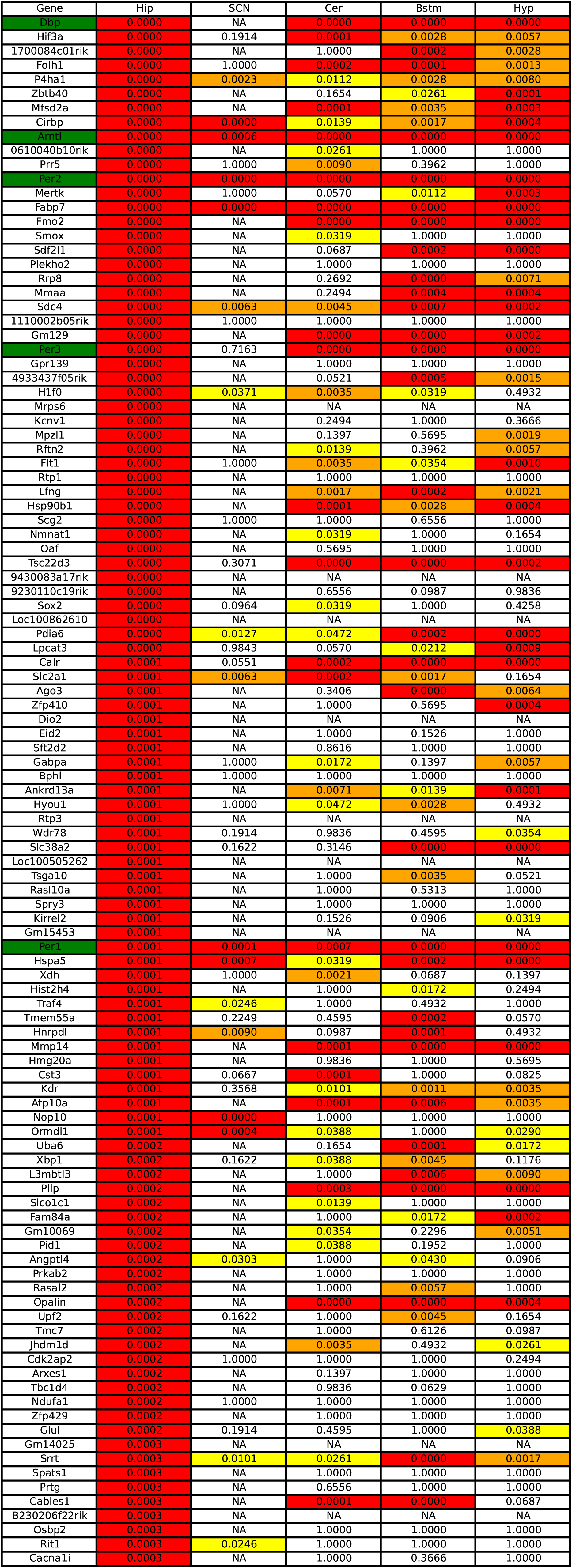
List of genes showing a circadian regulation in control mice, in the hippocampus (Hip), suprachiasmatic nucleus (SCN), Cerebellum (Cer), Brain Stem (Bstm) and Hypothalamus (Hyp). If most core clock genes display circadian regulation in most structures, numerous genes oscillate specifically in the hippocampus.

The analysis of the heatmaps (**Supplementary Fig. 2**) revealed that gene oscillations were distributed in a bimodal manner in control mice, with either a very low or very high oscillation amplitude (**Fig. 2b**). The majority of genes with high amplitude oscillations were phase advanced in comparison to the rest of the oscillatory transcripts (**Fig. 2b**).

Proteomic data enabled the analysis of 1,500 peptides. We found 170 peptides oscillating in controls (**Fig. 1e** and **Supplementary Table 2**), i.e. the same percentage (11%) as for gene transcripts. Nine proteins were oscillating in both proteomic and transcriptome datasets, which partly originates from the fact that 3% of peptides have a correspondence in the transcriptome dataset.

Together these results demonstrate that 10% of the gene/protein map in the hippocampus of control mice undergoes a continuous remapping during the night/day cycle. We then performed the same analysis in age-matched TLE mice in order to determine whether such circadian regulation was maintained in pathological conditions.

### Reprogramming of the circadian regulation of genes and proteins in experimental TLE

In experimental TLE, the number of oscillating transcripts was increased by 30% as compared to control (1650 versus 1256, **Fig. 1a** and **Supplementary Table 3**). Remarkably, only 29% of the oscillating transcripts were common to both control and TLE conditions, demonstrating a reprogramming of the daily remapping of hippocampal genes. Cluster analysis clearly separated the different groups by time and function (**Fig. 1b**). The core clock genes all gained in oscillation amplitudes in TLE as compared to control (**Fig. 1c,d**). Many important circadian-related genes gained oscillatory activity in TLE, from which we highlighted Bhlhe40 and Bhlhe41 (**Fig. 1d, 2a**). These genes encode proteins that interact with Arnlt and compete for E-box binding sites upstream of Per1 and repress Clock/Bmal activation of Per1, hence controlling circadian rhythms^25^. Increased oscillations of Bhlhe40 and Bhlhe41 may play a key role in the larger recruitment of oscillating transcripts in TLE. Although the mechanism of the reprogramming in TLE was outside the scope of this study, we identified a possible epigenetic mechanism. Runx1, a DNA binding regulator, gained statistical significance in TLE (**Fig. 2a**). Runx1 regulates Creb1, Cry1, Per1 and Per3, and has protein-protein interaction with two key histone deacetylases, Hdac1 and Hdac2. In keeping with this, proteomics revealed oscillatory activity for Hdac1 and Hdac2 in TLE (**Supplementary Table 4**). Finally, Hdac8 also gained oscillatory activity in TLE. Together, these results suggest the involvement of multiple epigenetic mechanisms in the reprogramming of the circadian regulation of hippocampal genes in TLE. The amplitudes of the oscillations were increased in TLE as compared to control, and genes with high amplitude oscillations were phase advanced as compared to the rest of the transcripts (**Fig. 2b** and **Supplementary Fig. 2**).

As for transcripts, the number of oscillating peptides was increased by 40% in TLE as compared to control (237 versus 170 peptides, Fig. 1E and table S4), with only 5% common to both experimental groups.

We conclude that the cycling of genes and proteins is very different in experimental TLE as compared to control, which makes comparing levels of expression, even at the same time point, more difficult.

### Functional analysis

The enriched terms reported from DAVID bioinformatics for oscillating genes found in control condition alone, epileptic condition alone, and oscillating in both conditions for three different p-values are listed in **Supplementary Tables 5-13**. Many biological pathways, including those linked to metabolism, showed circadian regulation in control and TLE, and specific alterations in TLE as revealed by KEGG pathways gene sets, Gene Ontology biological process, Gene Ontology metabolic function and REACTOME pathway database (**Supplementary Fig 3-14**, **Supplementary Table 14-25**).

**Figure 3.**
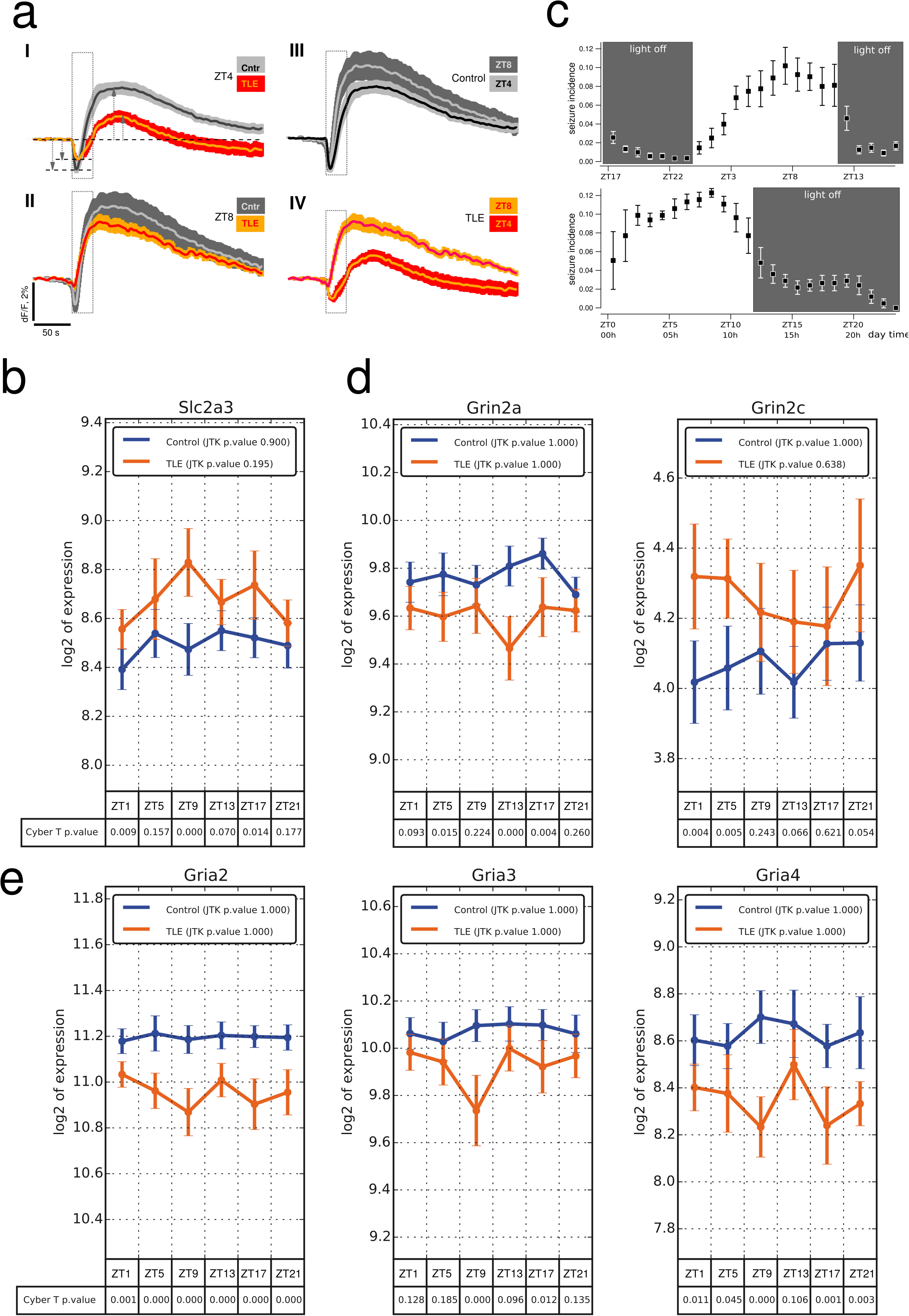
Alteration of circadian regulation of metabolism and drug targets in TLE. (**A**) (**a**) Averaged NAD(P)H fluorescence changes induced by 10 Hz 30s electrical pulse train stimulation of Schafer collaterals (rectangle) imaged in *Stratum Radiatum* of the hippocampal slices of control and TLE mice. (**a**) **I.** Averaged NAD(P)H fluorescence recorded in TLE (yellow trace) and control (dark gray trace) slices prepared at ZT4. The mean amplitude of the NAD(P)H transient overshoot was significantly smaller in TLE than in control mice (TLE: 1.3±0.17% n=9, control: 3.5±0.5%, n=10, p<0.01, dashed arrows) suggesting reduced activity of cytosolic glycolysis in TLE as compared with control. The decrease in NAD(P)H fluorescence dip amplitude in TLE did not reach statistical significance (TLE: −1.3±0.2%, control: −1.9±0.20%, p=0.2, dotted double-head arrows). **II.** There was no difference in NAD(P)H overshoot in slices prepared at ZT8 in TLE (red trace) and control (grey trace) mice (TLE: 4.1±0.4%, n=6, control: 4.6±0.9%, n=6, p=0.3). However, dip amplitudes were significantly different (TLE: −0.6±0.1%, control: −1.7±0.3%, p<0.05). A smaller dip indicates a reduced activity of oxidative phosphorylation or/and an increased activity of cytosolic glycolysis, which rapid onset produces a massive reduction of NAD^+^ and masks NADH oxidation in mitochondria. Tthe time of slice preparation had also a significant impact on metabolic activity in both controls (**III**) and TLE (**IV**) conditions. In slices made at ZT8 (grey trace) from control animals, NAD(P)H overshoot amplitude was slightly but significantly larger than in slice made at ZT4 (4.6±0.9%, n=6 vs 3.5±0.5%, n=10, p<0.05). The mean amplitudes of the initial dips were similar at both time points (−1.9±0.2% vs −1.7±0,4, p=0.7, n=6). In TLE slices made at ZT8 (red trace) NAD(P)H fluorescence rise started 8±2 s earlier (n>6, p<0.05) and reached greater levels than in slice made at ZT4 (yellow trace). (**b**) The type 3 glucose transporter shows a different circadian regulation in control and TLE, with a specific significant upregulation at ZT15 in TLE. (**c**) Top. Circadian regulation of seizure incidence during the night and day cycle. The highest seizure probability is found around ZT15. Bottom. Two days after shifting the light/dark cycle by 8 hours in the animal facility, the temporal pattern of seizure incidence shifted accordingly. (**d** and **e)** Alterations of the temporal expression of genes encoding for NMDA and AMPA receptors subunits, respectively.

In order to show how the resource we provide can be used to predict/test functional consequences, we focus, in this example, on metabolic pathways, which were altered in TLE. Metabolic activity was assessed *in vitro* in the ventral CA1 region of the hippocampus using NAD(P)H imaging in stratum radiatum during stimulation of Schaffer collaterals (**Fig. 3a**). Activation of hippocampal cells by Schaffer collateral stimulation results in characteristic changes in NAD(P)H fluorescence intensity: the initial ‘dip’ is followed by the ‘overshoot’ component both of which reflect enhanced oxidation and reduction of the pyridine nucleotides, respectively ^26^. In similar experimental conditions, the NAD(P)H fluorescence overshoot amplitude is mainly related to the intensity of cytosolic glycolysis in both astrocytes and neurons^26^. In slices prepared at ZT4, glycolysis was considerably decreased in TLE as compared to control (**Fig. 3a**), as previously reported ^27^. However, it was considerably increased in TLE in slices prepared at ZT8 (**Fig. 3a**), as compared to TLE at ZT4 and as compared to control at ZT8 (**Fig. 3a**). Interestingly, glycolysis was also different between ZT4 and ZT8 in control slices, suggesting a circadian control of metabolism in hippocampal neurons (**Fig. 3a**). In keeping with these results, there was a marked increase in the expression of the type 3 glucose transporter specifically at ZT8 in TLE (**Fig. 3b**). Hence, for a cellular process as fundamental as metabolism, different circadian functional modes characterize control and TLE conditions. Whether this is due to a reprogramming of the hippocampal clock remains to be determined.

The circadian rhythmicity can also be appreciated at the system level. Seizures are regulated in a circadian manner in human and experimental mesial TLE ^28^, a result we confirmed in our mouse model (**Fig. 3c**). One explanation could be that the daily remapping of the molecular landscape brings neuronal networks close to seizure threshold at specific time points^29^. In line with this scenario, we found different seizure susceptibility at different time points between control and TLE animals (**Supplementary Fig. 15**). This further exemplifies the fact that control and TLE animals are subject to different operating modes.

### Consequences for antiepileptic drug targets and chronotherapy

Another possible use of such resource is for drug target design and chronotherapy. Many common drugs (e.g. for gastritis, chronic pain, asthma, and rheumatoid arthritis) target products of genes characterized by circadian rhythmicity ^21^. We obtained information on drugs and genes from DrugBank database version 5.0.6 ^30^, which contains information about 8283 drugs including 62 drugs that exert antiepileptic and anticonvulsant effects. In the database, we found mammalian genes encoding 2230 drug targets, 35 drug carriers, 238 drug enzymes and 130 drug transporters. For the 62 antiepileptic/anticonvulsant drugs, we found genes encoding 102 drug targets, 1 drug carrier, 32 drug enzymes and 18 drug transporters.

Among the genes that oscillate in control and TLE animals (JTK_CYCLE adjusted p < 0.05), we found 24 and 71 drug targets, 2 and 2 drug carriers, 2 and 7 drug enzymes, and 2 and 9 drug transporters, respectively (**Table 2**). There was little overlap between control and experimental TLE datasets, further highlighting the necessity to take into account the reorganization of gene variations that is specific to experimental TLE. We obtained similar conclusions when looking at genes which expression changes (one-way ANOVA, FDR < 0.05) in at least one time point in control and TLE animals. We found 20 and 62 drug targets, 2 and 0 drug carriers, 2 and 11 drug enzymes, and 2 and 7 drug transporters, respectively (**Table 2**). **Table 3** provides examples of oscillating and differentially expressed gene products controlled by antiepileptic and anticonvulsant drugs.

**Table 2:** List of genes that oscillate (JTK_CYCLE adjusted p < 0.05) or that show changes in expression in at least one time point (one-way ANOVA, FDR < 0.05) in control animals and experimental TLE. Genes are separated in drug targets, carriers, enzymes and transporters. There is little overlap between the control and TLE conditions (common ones are underlined). The ABCC1 is not only transporter for *Phenobarbital* but also a drug target for Sulfinpyrazone and Biricodar dicitrate (in italics). Genes in Bold are shown in table 3.

| Type of action | Oscillating genes Control | Oscillating genes TLE | Expression change Control | Expression change TLE |
| --- | --- | --- | --- | --- |
| Target | ADAM10, BPHL, **CACNA1I**, CALR, CRBN, FLT1, FOLH1, <u>GLUL</u>, HSP90B1, HSPA5, IGF1R, KDR, <u>MERTK</u>, <u>MMAA</u>, <u>MMP14</u>, NDUFA1, NDUFA6, NMNAT1, NPEPPS, P4HA1, PRKAB2, <u>RAB8B</u>, SMOX, XDH | **ABCC1**, ACAN, ADRBK1, ADSSL1, AFG3L2, AKT2, ALAS1, ALDH6A1, ALKBH3, ANO1, APRT, BCL2, CD52, CTSK, CYB5R3, CYSLTR2, DDAH2, DHPS, DNPEP, EDNRB, F2R, FASN, FPGS, GARS, GLS, <u>GLUL</u>, GPT2, GRIK5, GRM7, GSR, HCN2, HDAC6, HDAC8, HIF1AN, HRH3, IMPG2, LARS, MAPT, <u>MERTK</u>, <u>MLLT4</u>, <u>MMAA</u>, <u>MMP14</u>, MTRR, NDUFA10, NFKBIA, PDE7A, PFN1, PIK3CA, PRKAB1, PSMA4, PSMA7, PSMB7, PTPRS, QPCT, <u>RAB8B</u>, RNGTT, RPL19, RPS6KA4, RYR2, SGPL1, SHMT1, SLC15A2, SLC16A3, SLC6A9, SLC7A8, STMN4, SULT2B1, TSTA3, TUBD1, VARS, WEE1 | ADAM10, BRAF, <u>CALR</u>, CXCR4, FLT1, FOLH1, <u>GLUL</u>, <u>HSP90B1</u>, <u>HSPA5</u>, <u>MERTK</u>, <u>MMAA</u>, NDUFA1, NDUFA6, <u>NFKBIA</u>, <u>P4HA1</u>, PRKAB2, SLC16A1, SMOX, TOP1, XDH | AASS, **ABCC1**, ADSSL1, AGT, AKT2, ALDH6A1, ALDH7A1, AMT, APRT, ASPH, CACNA1B, <u>CALR</u>, CD44, CYB5R3, CYSLTR2, DBH, DDAH2, DDX6, DHPS, DHRS3, DNPEP, F2R, **GABRA3**, <u>GLUL</u>, GPT2, GSR, GSTM5, HDAC6, HDAC8, HIF1AN, <u>HSP90B1</u>, <u>HSPA5</u>, HSPB1, IMPG2, KCNA4, LARS, **MAOB**, <u>MERTK</u>, <u>MLLT4</u>, <u>MMAA</u>, NCOA5, NDUFA10, <u>NFKBIA</u>, <u>P4HA1</u>, PAICS, PDE7A, PDK4, PRKAB1, PRODH, PSMA7, PTPRS, QPCT, RB1, RNGTT, ROCK2, SERPINF2, SHMT1, SLC7A8, SULT2B1, TUBD1, VARS, WEE1 |
| Carrier | FABP7, <u>SLC2A1</u> | FABP5, <u>SLC2A1</u> | FABP7, SLC2A1 |  |
| Enzyme | GLUG, XDH | APRT, FPGS, GLS, <u>GLUL</u>, GSR, HIF1AN, VARS | <u>GLUL</u>, XDH | APRT, CHKA, DBH, <u>GLUL</u>, GSR, HIF1AN, MAOB, PRODH, SRM, TGM1, VARS |
| Transporter | <u>SLC2A1</u>, **SLCO1C1** | **ABCC1**, ABCC5, SLC15A2, SLC16A3, <u>SLC2A1</u>, SLC2A9, SLC7A5, SLC7A8, TFRC | **SLC16A1**, SLC2A1 | **ABCC1**, ABCC5, SLC14A1, SLC2A9, SLC7A5, SLC7A8, TFRC |

**Table 3:** Genes that are drug target (T) or transporters (Tr), and that oscillate (JTK_CYCLE, adjusted p value < 0.05; OSC) or that are differentially expressed (one-way-ANOVA, adjusted p value < 0.05; DE) in control (CTR) or TLE animals. Information on drugs can be found in the Supplemental Material.

| Gene symbol | Function | OSC | DE | Animals | Anticonvulsant drug names |
| --- | --- | --- | --- | --- | --- |
| ABCC1 | Tr | + | + | TLE | Phenobarbital |
| CACNA1B | T | - | + | TLE | Gabapentin, levetiracetam |
| CACNA1I | T | + | - | CTR | Paramethadione, zonisamide, flunarizine |
| GABRA3 | T | - | + | TLE | Meprobamate, metharbital, thiopental, primidone, diazepam, methylphenobarbital, estazolam, fludiazepam, nitrazepam |
| MAOB | T | - | + | TLE | Zonisamide |
| SLC16A1 | Tr | - | + | CTR | Valproic Acid |
| SLCO1C1 | Tr | + | - | CTR | Phenytoin |

Finally, new classes of antiepileptic drugs targeting AMPA and NMDA receptors are being developed ^31^. Remarkably, many of the corresponding genes of AMPA and NMDA subunits present in the database showed different expression patterns at specific time points in TLE as compared to control (**Fig. 3d, e**).

Together these results suggest that the efficacy of antiepileptic and anticonvulsant drugs is time-dependent. It is important to stress that drug efficacy testing must be performed in experimental epilepsy models as the time-dependent regulation of the drug targets are different in TLE and control conditions. The resource we provide can be used to assess such factor.

## DISCUSSION

We conclude that hippocampal networks undergo a complex daily remapping at the gene and protein level. The number of oscillating genes and proteins is likely to be an underestimate as hippocampal samples were collected every four hours. The remapping may drive hippocampal networks into different functioning modes in a circadian-dependent manner to optimize information processing at specific time points. For example, the remapping may contribute to the circadian regulation of theta oscillations and place cell firing in the hippocampus ^32, 33^, or, if a remapping also occurs in the cortex, to the circadian regulation of cortical excitability in humans^34^. Such oscillations could be a downstream consequence of those occurring in the SCN, and the daily activation of various hormonal and neuromodulator pathways ^8^. Alternatively, the clock could be intrinsic since some hippocampal genes continue to oscillate *ex vivo* ^35^. Yet, the daily molecular remapping is important to consider for understanding hippocampal function. Since multiple other damped (secondary) circadian oscillators exist in the brain, each with specific phase properties ^36^, it is likely that a remapping also exists in other brain regions, each with a specific set of regulated genes and proteins. In addition, a similar resource should be provided for other hippocampal regions (like the dorsal hippocampus), since the molecular architecture of the hippocampus is not homogenous along its septo-temporal axis ^22^.

One major limitation of our approach is to lump all cell types from the different hippocampal subfields. Future studies should focus on cell-type specific analysis ^37^, a complex endeavor since it requires instantaneous “freezing” of cells and networks to stop the cycling process at specific times during the night and day cycle.

In epilepsy, we report a large reprogramming of the cycling behavior, with the possible involvement of epigenetic mechanisms ^38, 39^. It is not yet possible to determine whether this is a cause or consequence of seizures, altered sleep/wake cycles, or just a homeostatic mechanism. However, the important result is that the hippocampus uses a different functional regime in TLE as compared to control. This makes comparing genes and proteins from control and TLE tissues very difficult, as down or up-regulation could just sign different oscillating properties in both conditions (e.g. a phase shift). In fact, if two systems are under two different functioning modes, one can argue that they will have to be studied for themselves, rather than being compared to each other. Based on this novel information, it will be now important to indicate the circadian time a given animal was used in any experimental condition, and to take into consideration a possible reprogramming when studying neurological disorders. Indeed, we can propose that such reprogramming may occur in most (if not all) neurological disorders ^40^. This argument is even more crucial for chronotherapeutics ^20^, highlighting the necessity to take into account the fact that the expression of drug targets may display different expression time-dependent changes in pathologies as compared to controls.

The time of the day is thus a critical parameter to consider for the proper interpretation of molecular, physiological and behavioral studies, as the functional architecture of the hippocampus may be dynamically regulated in a time-dependent manner. Overall, the present findings have important implications for disease pathogenesis and therapy.

## Acknowledgments

This work was supported by Inserm, and the European Union’s Seventh Framework Programme (FP7/2007-2013) under grant agreement no602102 (EPITARGET). WL and SB are supported by the Niedersachsen-Research Network on Neuroinfectiology (N-RENNT) of the Ministry of Science and Culture of Lower Saxony in Germany. KJD and KL are supported by the Polish Ministry of Science and Higher Education grants 888/N-ESF-EuroEPINOMICS/10/2011/0 and W19/7.PR/2014.

## Note

All the data will be made publicly available on the Circadiomics ^1, 2^ web portal at http://circadiomics.igb.uci.edu/ upon acceptance of the manuscript.

## Supplementary Materials

### Materials and methods

#### Animals

All experiments were performed on FVB adult male mice following Inserm procedures. Twenty four control and twenty four TLE mice were used for transcriptomics and proteomics. They were kept in a special in house animal facility with a strict control of light and temperature conditions (beginning of the light phase at 7:30 and beginning of the night phase at 19:30). A red light, which does not disrupt circadian rhythmicity, was present during the night phase to allow researchers to manipulate the animals. During the night phase, no external light could enter the room when opening the door. Mice were housed in groups of four to five to enable social interaction. Cages had enriched environment. Throughout the experimental procedure, the same researcher took care of the animals at the same time of the day to limit external stressful factors. Animals were anesthetized with isoflurane in the animal facility and four of them killed every four hours (six time points).

The brain was quickly extracted and the hippocampus removed in modified ice cold ACSF. Right (for transcriptome analysis) and left (for proteomics) hippocampi were separated into their ventral and dorsal parts and quickly frozen. Only the ventral parts were used here. The average time between decapitation and sample freezing was 90s per mice to limit any degradation of gene products and proteins. All tissue collection was performed during a single 24h period. The collection of the hippocampal tissue from the four mice per time window was performed in less than 10 minutes.

#### Model of epilepsy

Adult FVB mice were injected with methylscopolamine (1 mg/kg i.p.) 30 min before the pilocarpine injections. Pilocarpine was repeatedly injected (100 mg/kg i.p.), every twenty minutes, until status epilepticus (SE) was observed ^3^. After 90 min of SE, we injected diazepam (10 mg/kg i.p.) to stop SE. All mice then received 0.5 ml NaCl (0.9%) subcutaneously, and again in the evening. During the following days, if required, mice were fed with a syringe.

#### 24/7 EEG monitoring

Ten mice were implanted with a telemetry probe (ETA F10, (Data Science International [DSI], St Paul, MN). The stereotaxic surgery was performed under ketamine (100mg/kg) / xylazine (10mg/kg) anesthesia. Lidocaine was used locally. Temperature, heart rate, and breathing rate were continuously monitored during the surgical procedure (MouseOx®Plus). A recording skull screw was secured above the hippocampal CA1 region (−1.8mm posterior, +1.8mm lateral), and a reference skull screw was secured above the cerebellum. The leads of the telemetry probe (Data Science International [DSI], St Paul, MN) were wrapped around the recording and reference screws, and screws were all encased in dental cement. The probe body containing the battery was placed subcutaneously in the back of the mice. Animals were allowed 7 days postsurgical recovery before any further experimental procedures were conducted. Electroencephalographic (EEG) recordings were performed continuously until the end of the experimental protocol. Seizures were automatically detected with DSI software (adjusting for frequency and amplitude) using low detection values. This led to numerous false positives (mostly movement artifacts), which were manually removed after double-checking by a trained technician and clinical epileptologist. Complete EEG recordings were fully visually inspected. No false negatives were detected (i.e., the semi-automated procedure insured the detection of all seizures). All pilocarpine-treated mice developed spontaneous seizures.

#### Animal care and handling

We kept 6 mice (control or epileptic) per cage, with enriched environment. We found that keeping epileptic mice together (as opposed to one mouse per cage) dramatically changed their behavior. They recovered faster from status epilepticus. They could be easily handled, denoting a lack of stress. When epileptic mice are housed in individual cages, they are extremely nervous and it is very difficult to handle them. Here, we removed a confounding stress factor linked to the lack of social interaction, as reported in rats ^4^.

#### Seizure threshold tests with pentylenetetrazole (PTZ)

Seizure threshold was determined in individual male FVB mice by the timed intravenous pentylenetetrazole (PTZ) test as described previously ^5^, in a total of 57 naive and 55 epileptic mice. Epilepsy was induced by pilocarpine ^3^, and seizure thresholds were determined 7-9 weeks after SE, when all mice had developed epilepsy. Groups of 8–20 mice were used per threshold determination. PTZ infusion was stopped at the first seizure, and seizure threshold was calculated in mg PTZ per kg body weight based on the infusion time needed to induce this seizure endpoint ^5^. PTZ thresholds were determined every 4 hours during the light and dark phase. The maximum number of threshold determinations was limited to 3 per animal with 7 days in between (repeated seizure threshold tests do not alter the threshold if an interval of at least 48 h lies within the determinations) ^5–7^. Additionally, we performed two PTZ seizure threshold determinations at the same time-point (ZT11:30) to compare PTZ thresholds at the beginning and at the end of the experiment, demonstrating the reproducibility of the method. Data were analyzed by using the Prism5 software from GraphPad (La Jolla, CA, USA). Thresholds of control and SE animals were analyzed separately using one-way analysis of variance (ANOVA) and Bonferroni post tests and combined by using a two-way ANOVA with Student’s *t*-tests as post hoc analysis. All tests were used two-sided; a *P* < 0.05 was considered significant.

#### RNA isolation and microarray hybridization

RNA isolation and microarray hybridization was performed as previously described ^8^. The isolation of total RNA was performed using the miRNeasy Mini kit (QIAGEN, # 217004) according to the manufacturer’s instructions. The sample quality was determined using a NanoDrop 2000 spectrophotometer (Thermo, Fisher Scientific) and Agilent 2100 Bioanalyzer. GeneChip® Mouse Gene 2.1 ST arrays (Affymetrix, Santa Clara, CA, # 902120) were used for mRNA profiling. One hundred nanograms of total mRNA was used for cDNA synthesis using the Ambion WT Expression Kit (Life Technologies, # 4411974). Hybridization, washing and scanning were conducted according to Affymetrix guidelines for the GeneAtlas^TM^ instrument.

#### Quantitative PCR

Reverse transcription for individual qPCRs was carried out using 250ng of total RNA and the High-Capacity Reverse Transcription Kit (# 4368814, Life Technologies) according to the manufacturer’s instruction. Quantitative PCRs were run on the ABI 79OOHT Fast Real-Time PCR instrument (Applied Biosystems). Gene expression analysis was conducted using a relative standard curve method with glyceraldehyde-3-phosphate dehydrogenase (Gapdh) to normalise the expression levels of target genes. All reactions were performed in triplicates.

#### Bioinformatic analysis

Analysis of microarrays was performed using R/Bioconductor (R Core Team, 2014; www.bioconductor.org). All microarrays were normalized with the Robust Multi-array Average (RMA) algorithm using oligo package (version 1.28.3) ^9^. Background value of intensity of probes was defined as median value of the intensity of antigenomic probes. The intensity of genomic probes below the background intensity was corrected to this background value. Probes with intensity greater than background value in less than four samples were removed from analysis. Only probes that corresponded to a single gene were selected for further analysis.

JTK_Cycle and BIO_CYCLE were used to identify oscillating genes at three p-value thresholds ^10^. Gene lists for transcripts oscillating at each p-value in the control condition alone, the TLE condition alone, or both conditions were identified for further analysis. Cyber-T was used to identify differentially expressed timepoints between control and TLE conditions for each animal ^11^. Cyber-T p-values found for each time point for core clock genes are reported (Figure 2C). Proteomic data for both control and TLE conditions was also analyzed by JTK_Cycle to identify rhythmic proteins. All transcriptome data is available for search on Circadiomics along with JTK_Cycle and BIO_CYCLE results ^1^.

Genes, which expression level differed between time points according to one-way ANOVA analysis performed separately for control and epileptic animals are presented on the heatmap (Figure 1). Differences were considered as significant for adjusted p-value < 0.5 – or 0.01 – depending on our final decision (FDR – Benjamini & Hochberg (BH) correction). Genes were ordered by clustering complete-linkage method together with Pearson correlation distance measure. Nine gene clusters highlighted on the heatmap were obtained by cutting dendrogram the selected level of the dendrogram. Functional analysis for Biological Function Go Terms was performed using MF FAT Chart option with default settings in DAVID with (http://david.abcc.ncifcrf.gov; ^12, 13^). Transcription factor binding sites overrepresented in promoters within clusters were detected with gProfiler (http://biit.cs.ut.ee/gprofiler/; ^14, 15^) with custom background defined as genes detected in the experiment to reveal the functional and regulatory groups presented in the data.

Further transcription factor binding site enrichment analysis was performed using MotifMap^16, 17^ and CHiP-seq datasets provided by UCSC Encode project ^18^. Transcription factor binding sites identified by MotifMap were found both 10 KB upstream and 2 KB downstream of transcription start sites (TSS) for all genes in each of the differentially expressed gene lists for all pair-wise comparisons. The motifs identified were found with a BBLS conservation score of greater than or equal to 1.0 and an FDR of less than 0.25. CHiP-seq datasets were used to locate experimentally identified binding site locations within the same interval around the TSS for differentially expressed genes. Only the 90th percentile of CHiP-seq peaks were used from each dataset provided by UCSC Encode. A Fisher’s exact test was performed separately using the full background list of motifs from MotifMap and all target genes from the CHiP-seq datasets to compute the statistical likelihood of over representation in promotor regions.

### Proteomics

#### Protein extraction

All steps of the protein extraction were performed on ice or at 4°C. Isolated hippocampal tissue was homogenized in lysis buffer containing 20 mM triethylammonium bicarbonate (TEAB, Fluka), 5 % sodium deoxycholate (SDC, Sigma-Aldrich) and 1 tablet of protease inhibitor (Roche). Homogenates were boiled for 3 minutes and subsequently sonicated three times for 10 s, each time paused for 30 s.

Samples were centrifuged to pellet tissue debris and the protein concentration in the supernatant was measured with a BCA protein assay (Pierce) using bovine serum albumin as standard.

#### Protein Digestion

Dithiotreitol (DTT) was added to 200 µg of the global protein fraction up to a final concentration of 20 mM DTT in the solution. Samples were incubated for 20 min at 55°C and then centrifuged for 1 minute at 10000xg (20 °C). The supernatant was transferred to filter devices (10kDa MWCO, VWR) and filled up with digestion buffer (DB) containing 20 mM TEAB and 0.5 % SDC. Next, the samples were centrifuged for 10 minutes at 10000xg (20°C), the flow-through was discarded and 100 µl of 40 mM Indoacetamide (IAA) in DB was added to the filter devices followed by a 20 min incubation at room temperature in darkness. Afterwards samples were washed twice with DB and trypsin solved in DB was added to yield a protein to trypsin ratio of 100:1. Samples were incubated over night at 37°C. Filter devices were centrifuged for 20 minutes at 10000xg (20° C) and the flow-through was collected. Residual peptides were removed by another centrifugation step with DB. The flow-throughs of one sample were combined and first an equal volume of ethyl acetate and subsequently trifluoreoacetic acid (TFA, final concentration 0.5%) was added. The solution was mixed and centrifuged for 2 minutes at 14000xg (20° C). The upper layer was discarded and 250 µl of acetate ethyl was added, followed by centrifugation. The upper layer was discarded again and the aqueous phase was collected for further procession.

#### Isobaric labeling and peptide fractionation

Peptides were vacuum concentrated and labeled with amine-reactive, 6-plex tandem mass tag reagents (Thermo Fisher Scientific, Bremen, Germany) according to manufacturer’s instructions. The labeling reaction was quenched by addition of 5% hydroxylamine. Labeled peptides were pooled and desalted on Oasis HLB cartridges (Waters GmbH, Eschborn, Germany). Eluates containing 70% acetonitrile, 0.1% formic acid (FA) were dried and fractionated to 24 fractions by isoelectric point with an Offgel fractionator according to manufacturer’s recommendations (Agilent Technologies, Waldbronn, Germany). Peptide fractions were dried and stored at −20°C.

### LC-MS analysis

Peptides were dissolved in 8 µl 0.1% trifluoroacetic acid. 1.5 µl were injected onto a C18 trap column (20 mm length, 100 µm inner diameter) coupled to a C18 analytical column (200 mm length, 75 µm inner diameter), made in house with 1.9 µm ReproSil-Pur 120 C18-AQ particles (Dr. Maisch, Ammerbuch, Germany). Solvent A was 0.1% formic acid. Peptides were separated during a linear gradient from 4% to 40% solvent B (90% ACN, 0.1% formic acid) within 90 min. The nanoHPLC was coupled online to an LTQ Orbitrap Velos mass spectrometer (Thermo Fisher Scientific). Peptide ions between 330 and 1800 m/z were scanned in the Orbitrap detector with a resolution of 30,000 (maximum fill time 400 ms, AGC target 10^6^, lock mass 371.0318 Da). The 20 most intense precursor ions (threshold intensity 5000) were subjected to higher energy collision induced dissociation (HCD) and fragments also analyzed in the Orbitrap. Fragmented peptide ions were excluded from repeat analysis for 15 s. Raw data processing and analyses of database searches were performed with Proteome Discoverer software 1.4.1.12 (Thermo Fisher Scientific). Peptide identification was done with an in house Mascot server version 2.4.1 (Matrix Science Ltd, London, UK). MS2 data (including a-series ions) were searched against mouse sequences from SwissProt (release 2014_01). Precursor Ion m/z tolerance was 10 ppm, fragment ion tolerance 20 mmu. Tryptic peptides were searched with up to two missed cleavages. Low scoring spectrum matches were searched again with semitryptic specificity with up to one missed cleavage. Oxidation (Met), acetylation (protein N-terminus), and TMTsixplex (on Lys and N-terminus) were set as dynamic modifications, carbamidomethylation (Cys), was set as static modification. Mascot results from searches against SwissProt were sent to the percolator algorithm ^19^ version 2.04 as implemented in Proteome Discoverer. Only proteins with two peptides (maximum FDR 1%) were considered identified.

### *In vitro* experiments

#### Tissue slice preparation

Mice were anaesthetized with isoflurane and decapitated at ZT11 or ZT15. The brain was rapidly removed from the skull and placed in the ice-cold ACSF as above. The ACSF solution consisted of (in mmol/L): NaCl 126, KCl 3.50, NaH2PO4 1.25, NaHCO3 25, CaCl2 2.00, MgCl2 1.30, and dextrose 5, pH 7.4. ACSF was aerated with 95% O2/5% CO2 gas mixture. Slices of ventral hippocampus (350 µm) were cut as described ^20^, using a tissue slicer (Leica VT 1200s, Leica Microsystem, Germany). The ice-cold (< 6°C) cutting solution consisted of (in mmol/L): K-gluconate 140, HEPES 10, Na-gluconate 15, EGTA 0.2, NaCl 4, pH adjusted to 7.2 with KOH. Slices were then immediately transferred to a multi-section, dual-side perfusion holding chamber with constantly circulating ACSF and allowed to recover for 2h at room temperature (22°C-24°C). Slices were then transferred to a recording chamber continuously superfused with ACSF (flow rate 7ml/min, warmed at 30–31°C) with access to both slice sides.

#### Synaptic stimulation and field potential recordings

Schaffer collaterals were stimulated using a DS2A isolated stimulator (Digitimer Ltd, UK) with a bipolar metal electrode. Stimulus current was adjusted using single pulses (100-300 µA, 200µs, 0.15 Hz) to induce a local field potential (LFP) of about 60% of maximal amplitude. LFPs were recorded using glass microelectrodes filled with ASCF, placed in stratum pyramidale and connected to an EXT-02F/2 amplifier (NPI Electronic GmbH, Germany). Synaptic stimulation consisting of a stimulus train (200µs pulses) at 10 Hz lasting 30s was used to trigger metabolic response.

### NAD(P)H fluorescence imaging

NAD(P)H and NADH have similar optical properties, therefore it is expected that NAD(P)H may contribute to the total autofluorescence signal. Because the cellular NADP/ NAD(P)H pool is about one order of magnitude lower than the NAD/NADH pool, in the present study we assume that short-term variations of experimentally detected NAD(P)H responses account for variations in NADH (see discussion in ^21^). Changes in NAD(P)H fluorescence in hippocampal slices were monitored using a 290–370 nm excitation filter and a 420 nm long-pass filter for the emission (Omega Optical, Brattleboro, VT). The light source was the pE-2 illuminator (CoolLed, UK) equipped with 365 nm LED. Slices were epi-illuminated and imaged through a Nikon upright microscope (FN1, Eclipse) with 4x/0.10 Nikon Plan objective. Images were acquired using a 16-bit Pixelfly CCD camera (PCO AG, Germany). Because of a low level of fluorescence emission, NAD(P)H images were acquired every 600-800 ms as 4×4 binned images (effective spatial resolution of 348×260 pixels). The exposure time was adjusted to obtain baseline fluorescence intensity between 2000 and 3000 optical intensity levels. Fluorescence intensity changes in stratum radiatum near the site of LFP recording were measured in 3 regions of interest ∼1mm distant from the stimulation electrode tip using ImageJ software (NIH, USA). Data were expressed as the percentage changes in fluorescence over a baseline [(ΔF/F) • 100]. Signal analysis was performed using IgorPro software (WaveMetrics, Inc, OR, USA).

### Statistical analysis and signal processing

NAD(P)H dip and overshoot amplitudes were expressed as means ±SEM. Normality was rejected by Shapiro-Wilk normality test; therefore we used non-parametric Wilcoxon rank sum test. The level of significance was set at p<O.OS.

## Author contributions

AG, AMB GB and JAM performed experiments, and analyzed data. AB, AI, KJD, NG, SB, SJ, SS and WL performed experiments, analyzed data and wrote the manuscript. AB, KL, PB and PSC supervised aspects of the project and wrote the manuscript. CB conceived and managed the project, performed experiments, analyzed data and wrote the manuscript.

## Additional information about drugs

**Sulfinpyrazone** (DB01138) - A uricosuric drug that is used to reduce the serum urate levels in gout therapy. It lacks anti-inflammatory, analgesic, and diuretic properties. [PubChem] INDICATION: For the treatment of gout and gouty arthritis.

**Biricodar dicitrate** (DB04851) - The pipecolinate derivative biricodar (VX-710) is a clinically applicable modulator of P-glycoprotein (Pgp) and multidrug resistance protein (MRP-1). INDICATION: Administered intravenously, biricodar dicitrate is to be used in combination with cancer chemotherapy agents.

## Antiepileptic and Anticonvulsant drugs

**Diazepam** (DB00829) - A benzodiazepine with anticonvulsant, anxiolytic, sedative, muscle relaxant, and amnesic properties and a long duration of action. Its actions are mediated by enhancement of gamma-aminobutyric acid activity. It is used in the treatment of severe anxiety disorders, as a hypnotic in the short-term management of insomnia, as a sedative and premedicant, as an anticonvulsant, and in the management of alcohol withdrawal syndrome. (From Martindale, The Extra Pharmacopoeia, 30th ed, p589) INDICATION: Used in the treatment of severe anxiety disorders, as a hypnotic in the short-term management of insomnia, as a sedative and premedicant, as an anticonvulsant, and in the management of alcohol withdrawal syndrome.

**Estazolam** (DB01215) - A benzodiazepine with anticonvulsant, hypnotic, and muscle relaxant properties. It has been shown in some cases to be more potent than diazepam or nitrazepam. [PubChem] INDICATION: For the short-term management of insomnia characterized by difficulty in falling asleep, frequent nocturnal awakenings, and/or early morning awakenings.

**Fludiazepam** (DB01567) - Fludiazepam is a drug which is a benzodiazepine derivative. It possesses anxiolytic, anticonvulsant, sedative and skeletal muscle relaxant properties. It is a scheduled drug in the U.S., but is approved for use in Japan. INDICATION: Used for the short-term treatment of anxiety disorders.

**Flunarizine** (DB04841) - Flunarizine is a selective calcium entry blocker with calmodulin binding properties and histamine H1 blocking activity. It is effective in the prophylaxis of migraine, occlusive peripheral vascular disease, vertigo of central and peripheral origin, and as an adjuvant in the therapy of epilepsy. INDICATION: Used in the prophylaxis of migraine, occlusive peripheral vascular disease, vertigo of central and peripheral origin, and as an adjuvant in the therapy of epilepsy.

**Gabapentin** (DB00996) - Gabapentin (brand name Neurontin) is a medication originally developed for the treatment of epilepsy. Presently, gabapentin is widely used to relieve pain, especially neuropathic pain. Gabapentin is well tolerated in most patients, has a relatively mild side-effect profile, and passes through the body unmetabolized. INDICATION: For the management of postherpetic neuralgia in adults and as adjunctive therapy in the treatment of partial seizures with and without secondary generalization in patients over 12 years of age with epilepsy.

**Levetiracetam** (DB01202) - Levetiracetam is an anticonvulsant medication used to treat epilepsy. Levetiracetam may selectively prevent hypersynchronization of epileptiform burst firing and propagation of seizure activity. Levetiracetam binds to the synaptic vesicle protein SV2A, which is thought to be involved in the regulation of vesicle exocytosis. Although the molecular significance of levetiracetam binding to synaptic vesicle protein SV2A is not understood, levetiracetam and related analogs showed a rank order of affinity for SV2A which correlated with the potency of their antiseizure activity in audiogenic seizure-prone mice. INDICATION: Used as adjunctive therapy in the treatment of partial onset seizures in adults and children 4 years of age and older with epilepsy.

**Meprobamate** (DB00371) - A carbamate with hypnotic, sedative, and some muscle relaxant properties, although in therapeutic doses reduction of anxiety rather than a direct effect may be responsible for muscle relaxation. Meprobamate has been reported to have anticonvulsant actions against petit mal seizures, but not against grand mal seizures (which may be exacerbated). It is used in the treatment of anxiety disorders, and also for the short-term management of insomnia but has largely been superseded by the benzodiazepines. (From Martindale, The Extra Pharmacopoeia, 30th ed, p603) Meprobamate is a controlled substance in the U.S. INDICATION: For the management of anxiety disorders or for the short-term relief of the symptoms of anxiety.

**Metharbital** (DB00463) - Metharbital was patented in 1905 by Emil Fischer working for Merck. It was marketed as Gemonil by Abbott Laboratories. It is a barbiturate anticonvulsant, used in the treatment of epilepsy. It has similar properties to phenobarbital [Wikipedia]. INDICATION: Metharbital is used for the treatment of epilepsy.

**Methylphenobarbital** (DB00849) - A barbiturate that is metabolized to phenobarbital. It has been used for similar purposes, especially in epilepsy, but there is no evidence mephobarbital offers any advantage over phenobarbital. [PubChem] INDICATION: For the relief of anxiety, tension, and apprehension, also used as an anticonvulsant for the treatment of epilepsy.

**Nitrazepam** (DB01595) - A benzodiazepine derivative used as an anticonvulsant and hypnotic. INDICATION: Used to treat short-term sleeping problems (insomnia), such as difficulty falling asleep, frequent awakenings during the night, and early-morning awakening.

**Paramethadione** (DB00617) - Paramethadione is an anticonvulsant in the oxazolidinedione class. It is associated with fetal trimethadione syndrome, which is also known as paramethadione syndrome. INDICATION: Used for the control of absence (petit mal) seizures that are refractory to treatment with other medications.

**Phenobarbital** (DB01174) - A barbituric acid derivative that acts as a nonselective central nervous system depressant. It promotes binding to inhibitory gamma-aminobutyric acid subtype receptors, and modulates chloride currents through receptor channels. It also inhibits glutamate induced depolarizations. [PubChem]. INDICATION: For the treatment of all types of seizures except absence seizures.

**Phenytoin** (DB00252) - An anticonvulsant that is used in a wide variety of seizures. It is also an anti-arrhythmic and a muscle relaxant. The mechanism of therapeutic action is not clear, although several cellular actions have been described including effects on ion channels, active transport, and general membrane stabilization. The mechanism of its muscle relaxant effect appears to involve a reduction in the sensitivity of muscle spindles to stretch. Phenytoin has been proposed for several other therapeutic uses, but its use has been limited by its many adverse effects and interactions with other drugs. [PubChem]. INDICATION: For the control of generalized tonic-clonic (grand mal) and complex partial (psychomotor, temporal lobe) seizures and prevention and treatment of seizures occurring during or following neurosurgery.

**Primidone** (DB00794) - An antiepileptic agent related to the barbiturates; it is partly metabolized to phenobarbital in the body and owes some of its actions to this metabolite. Adverse effects are reported to be more frequent than with phenobarbital. (From Martindale, The Extra Pharmacopoeia, 30th ed, p309) INDICATION: For the treatment of epilepsy

**Thiopental** (DB00599) - A barbiturate that is administered intravenously for the induction of general anesthesia or for the production of complete anesthesia of short duration. It is also used for hypnosis and for the control of convulsive states. It has been used in neurosurgical patients to reduce increased intracranial pressure. It does not produce any excitation but has poor analgesic and muscle relaxant properties. Small doses have been shown to be anti- analgesic and lower the pain threshold. (From Martindale, The Extra Pharmacopoeia, 30th ed, p920) INDICATION: For use as the sole anesthetic agent for brief (15 minute) procedures, for induction of anesthesia prior to administration of other anesthetic agents, to supplement regional anesthesia, to provide hypnosis during balanced anesthesia with other agents for analgesia or muscle relaxation, for the control of convulsive states during or following inhalation anesthesia or local anesthesia, in neurosurgical patients with increased intracranial pressure, and for narcoanalysis and narcosynthesis in psychiatric disorders.

**Valproic Acid** (DB00313) - Valproic acid, supplied as the sodium salt valproate semisodium or divalproex sodium, is a fatty acid with anticonvulsant properties used in the treatment of epilepsy. The mechanisms of its therapeutic actions are not well understood. It may act by increasing gamma-aminobutyric acid levels in the brain or by altering the properties of voltage dependent sodium channels. Typically supplied in the sodium salt form (CAS number: 76584-70-8). Valproic Acid is also a histone deacetylase inhibitor and is under investigation for treatment of HIV and various cancers. INDICATION: For treatment and management of seizure disorders, mania, and prophylactic treatment of migraine headache. In epileptics, valproic acid is used to control absence seizures, tonic-clonic seizures (grand mal), complex partial seizures, and the seizures associated with Lennox-Gastaut syndrome.

**Zonisamide** (DB00909) - Zonisamide is a sulfonamide anticonvulsant approved for use as an adjunctive therapy in adults with partial-onset seizures. Zonisamide may be a carbonic anhydrase inhibitor although this is not one of the primary mechanisms of action. Zonisamide may act by blocking repetitive firing of voltage-gated sodium channels leading to a reduction of T-type calcium channel currents, or by binding allosterically to GABA receptors. This latter action may inhibit the uptake of the inhibitory neurotransmitter GABA while enhancing the uptake of the excitatory neurotransmitter glutamate. INDICATION: For use as adjunctive treatment of partial seizures in adults with epilepsy.

---

**figure S1.**
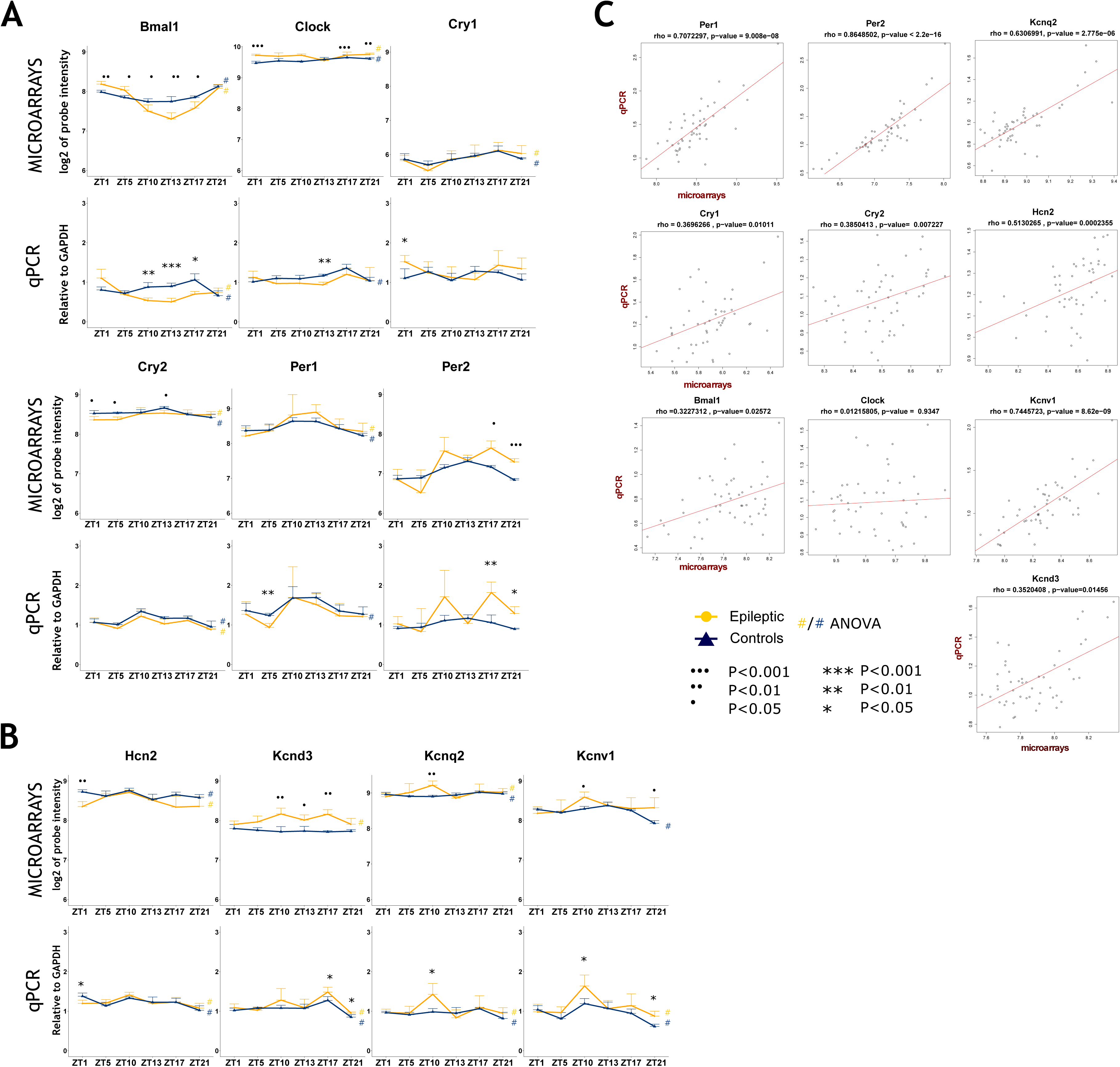
Changes in expression levels of selected genes in the ventral hippocampus of TLE and control animals. Panel A shows results for clock genes, panel B is dedicated for selected channels genes. Top rows show microarray data, whilst bottom rows show qPCR results. In the qPCR analysis GAPDH was used as an internal control and all values represent the mRNA level of the gene of interest normalised to the GAPDH mRNA level. Results are represented as means ± SD (n=4 in each group for each time-point in TLE and control animals). Note the correspondence between microarray and qPCR data. Panel C shows the correlation between qPCR and microarray data calculated using Spearman correlation. Rho and p-values are shown on the graph. Yellow circles represent epileptic animals and blue triangles represent control animals. Symbols indicate # results of the one-way ANOVA analysis, * p<0.05, ** p<0.01 of unpaired t-test with Welch’s correction, ooo p<0.001, oo p<0.01, o p<0.05.

**figure S2.**
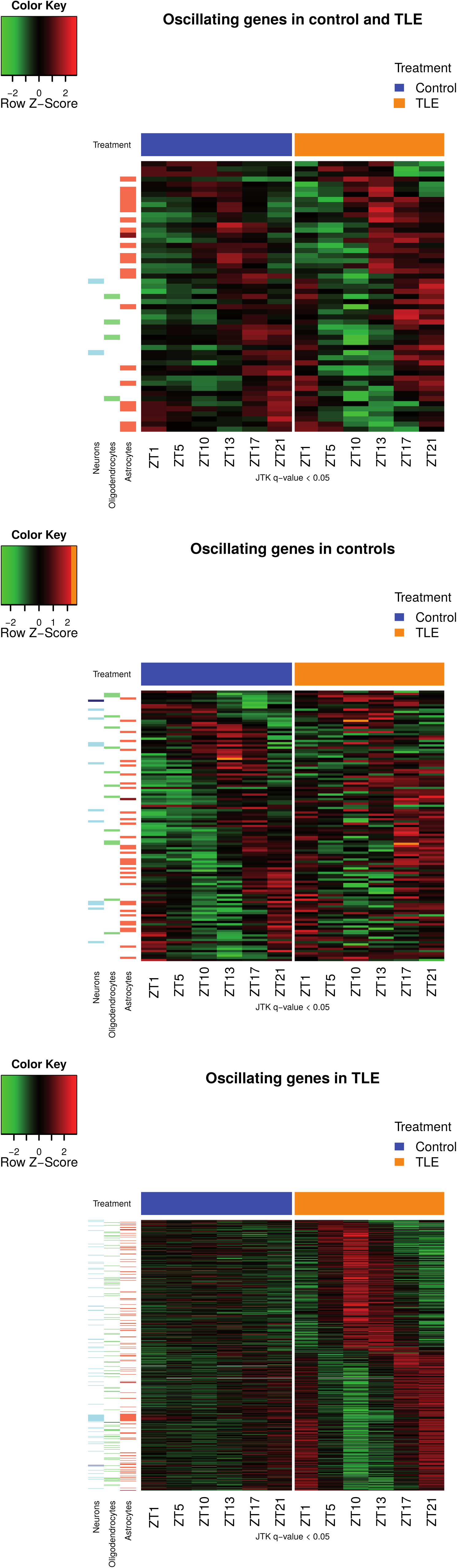
Heatmaps of oscillating genes in both conditions (top), control only (middle) and epileptic (TLE) only (bottom). JTK algorithm ^10^ was used to estimate gene oscillations. A gene was considered as significantly oscillating if Benjamani-Hochberg q-value was less then 0.05. Genes were ordered according to phase. The treatment groups are color-coded: blue for controls and orange for TLE. The panels on the left side of the heatmaps show which genes are enriched in neurons (blue), oligodendrocytes (green), and astrocytes (coral). Cell type enrichment is based on data obtained from ^22^.

**figures S3 to figure S8.**
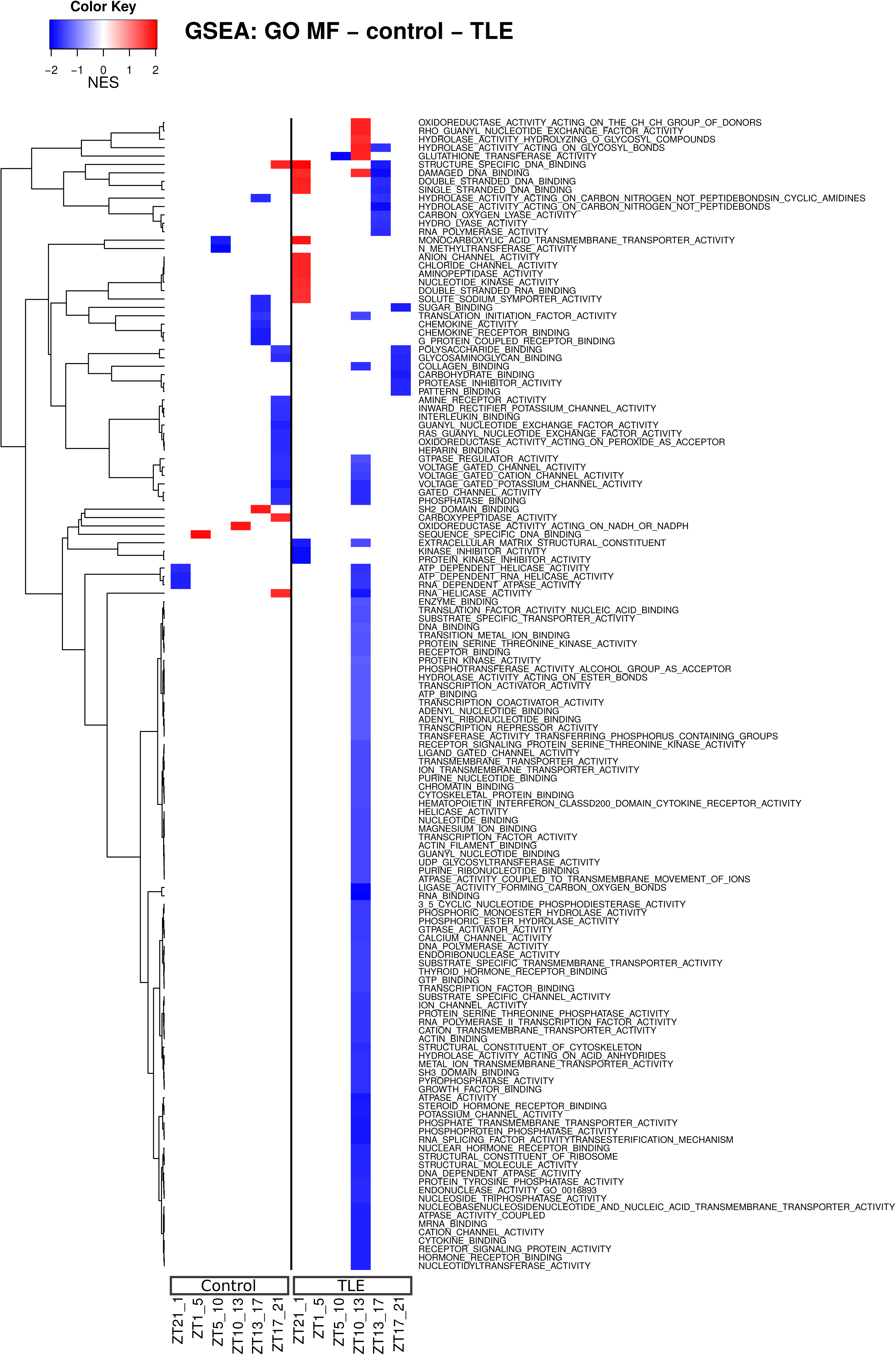
Differences in up- or down-regulation of gene sets for Gene Ontology Molecular Function (GO_MF, S3), Cellular Component (GO_CC, S4) and Biological Process (GO_BP, S5), as well as for biological pathways based on BIOCARTA (S6), KEGG (S7) and REACTOME (S8). Gene Sets were obtained from Molecular Signatures Database v5.0 (http://software.broadinstitute.org/gsea/msigdb/index.jsp) and used in Gene Set Enrichment Analysis (http://software.broadinstitute.org/gsea/index.jsp). For GSEA, we used a ranking based on score -log10(pvalue) * sign (fold change). We calculated p-values comparing two consecutive time points with t-test in control and epileptic animals separately (e.g. when comparing ZT3 and ZT7, we used ZT3 as our reference time point). All values can be found in corresponding tables S14 to S19. A large wealth of information is available to evaluate the circadian regulation of different pathways in controls and their modification in TLE. For example, based on GO_MF, there is a down-regulation of voltage-gated channels at ZT3 in controls, but at ZT19 in TLE; whilst anion channel activity is specifically up-regulated at ZT7 in TLE.

**figure s4.**
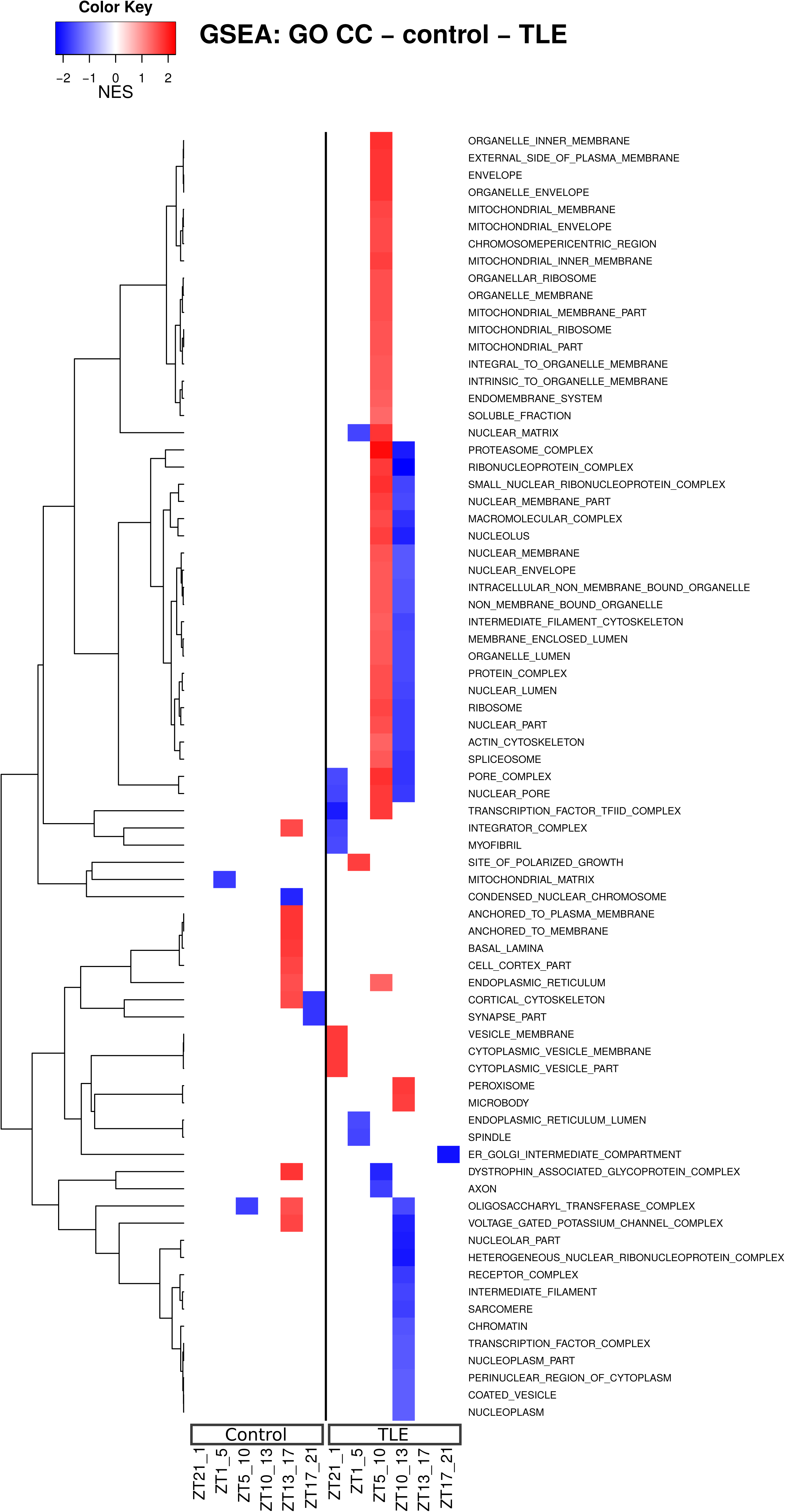

**figure s5.**
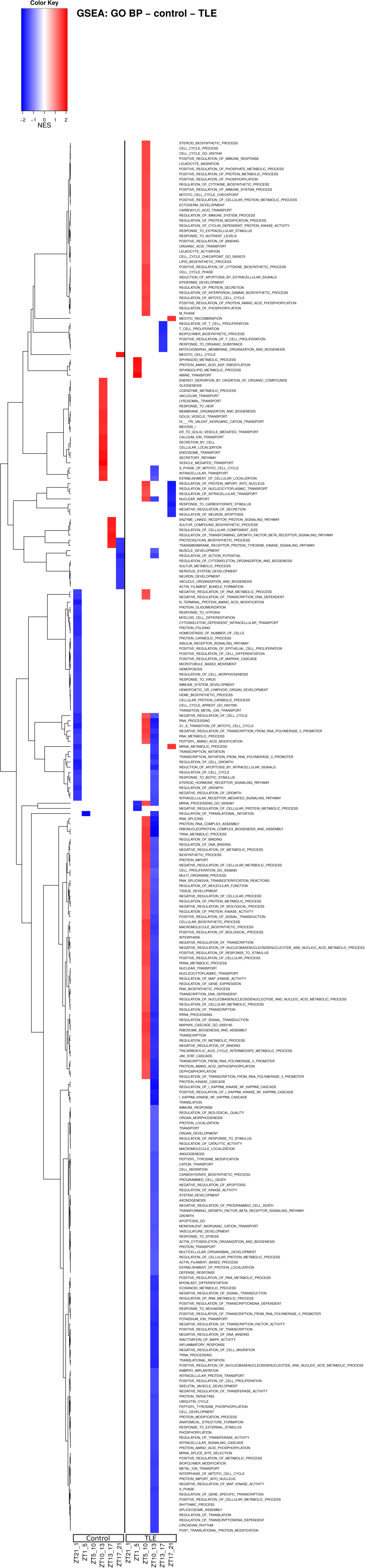

**figure s6.**
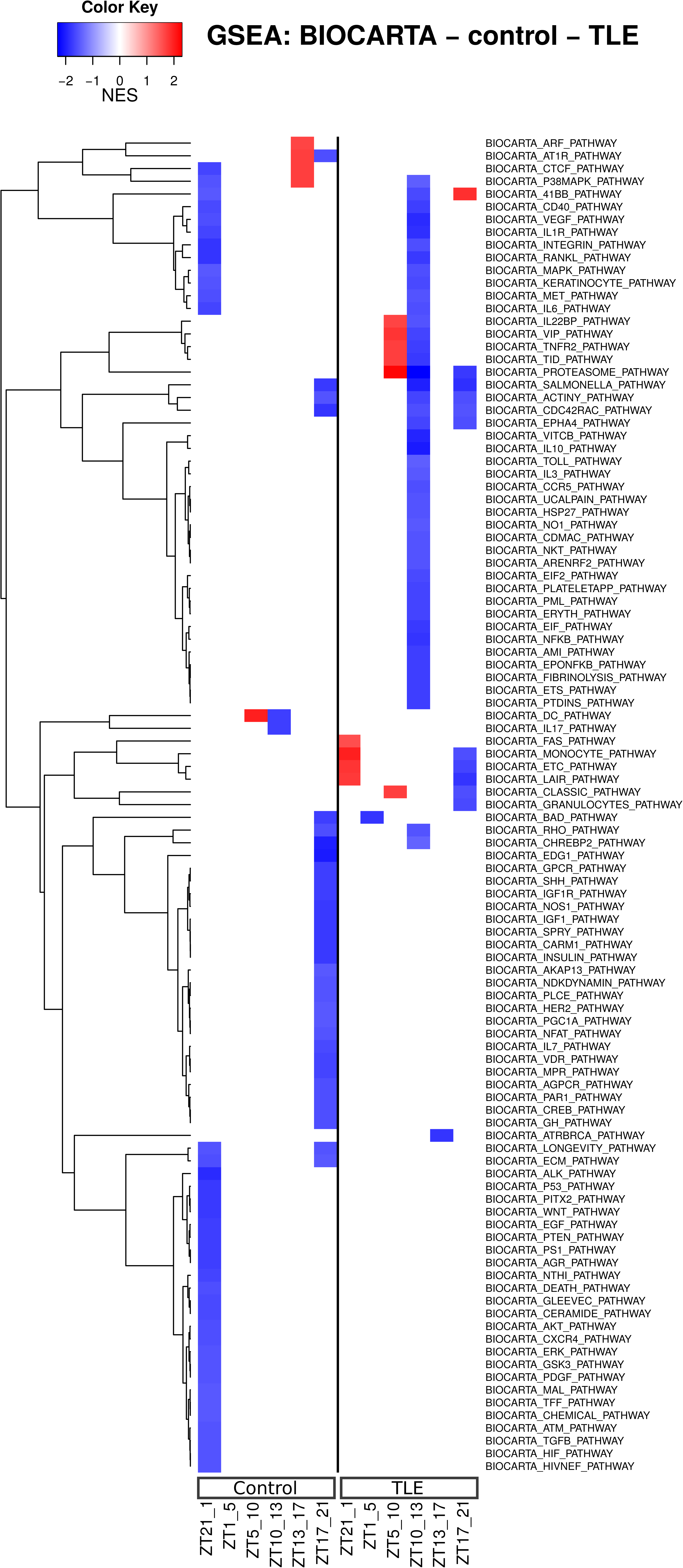

**figure s7.**
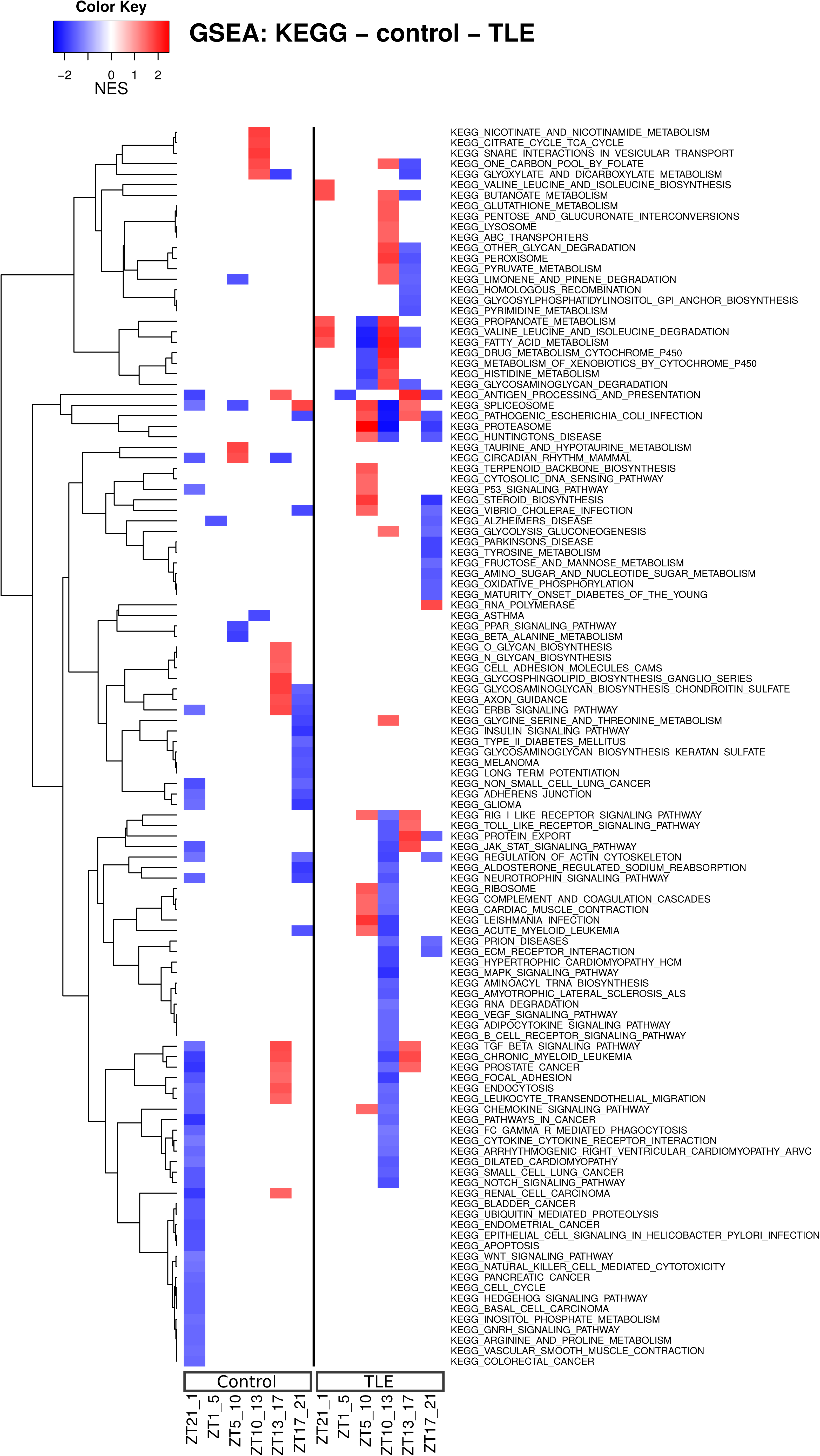

**figure s8.**
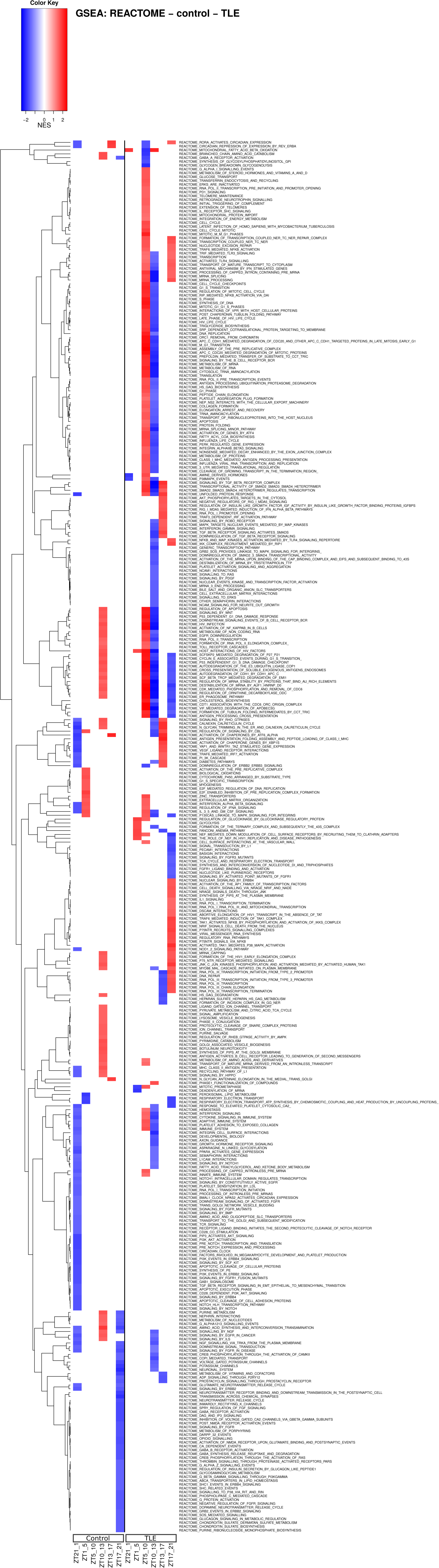

**figures S9 to figure S14.**
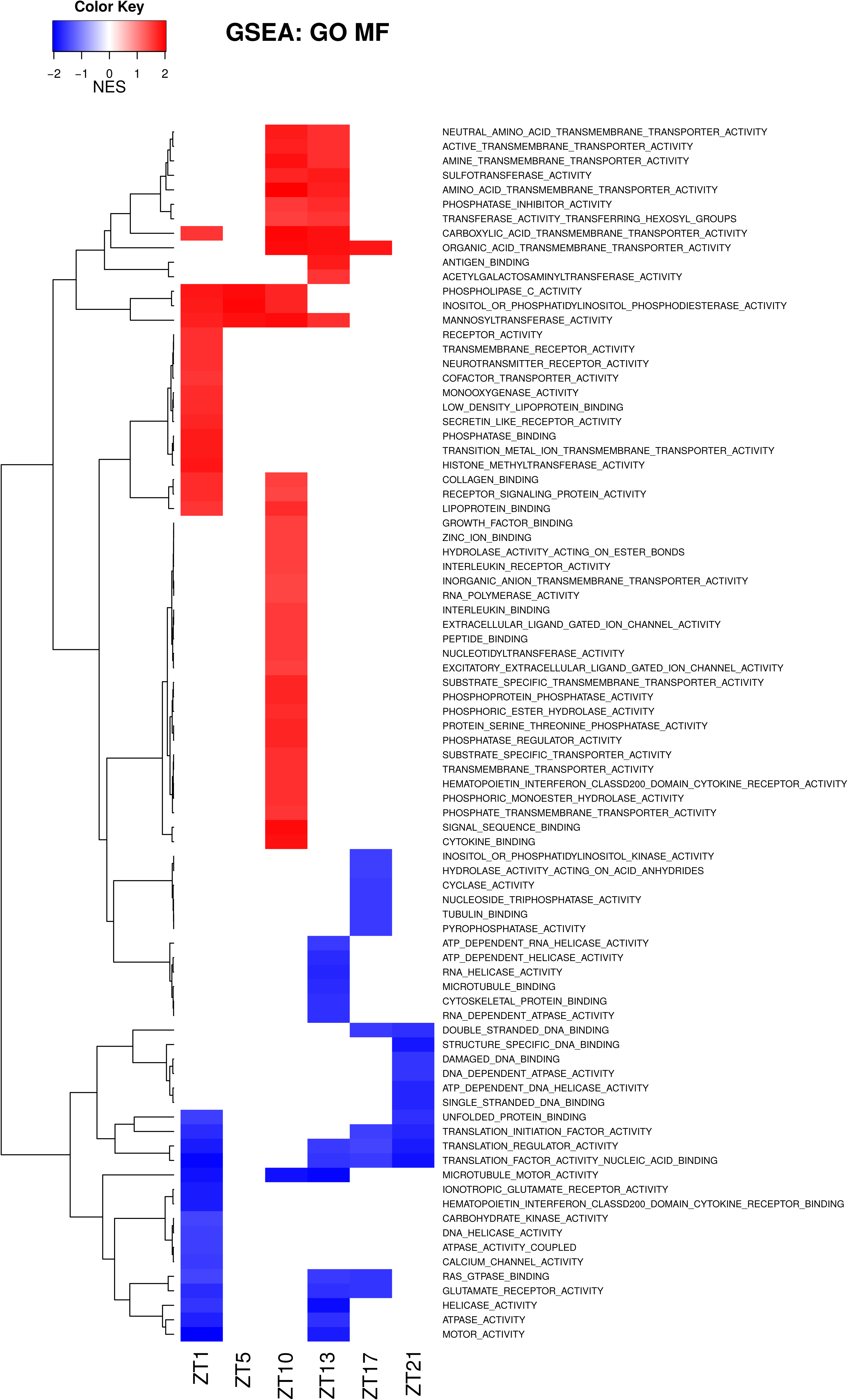
As figure S3 to S8, but here the analysis was done to compare differences between control and TLE animals at each time point. All values can be found in corresponding tables S20 to S25. This data is particularly useful to identify which pathways are modified in TLE at specific time points. For example, there is an up-regulation of glucose metabolism at ZT19 and a down-regulation of pyruvate metabolism at ZT3 and ZT7 in TLE.

**figure s10.**
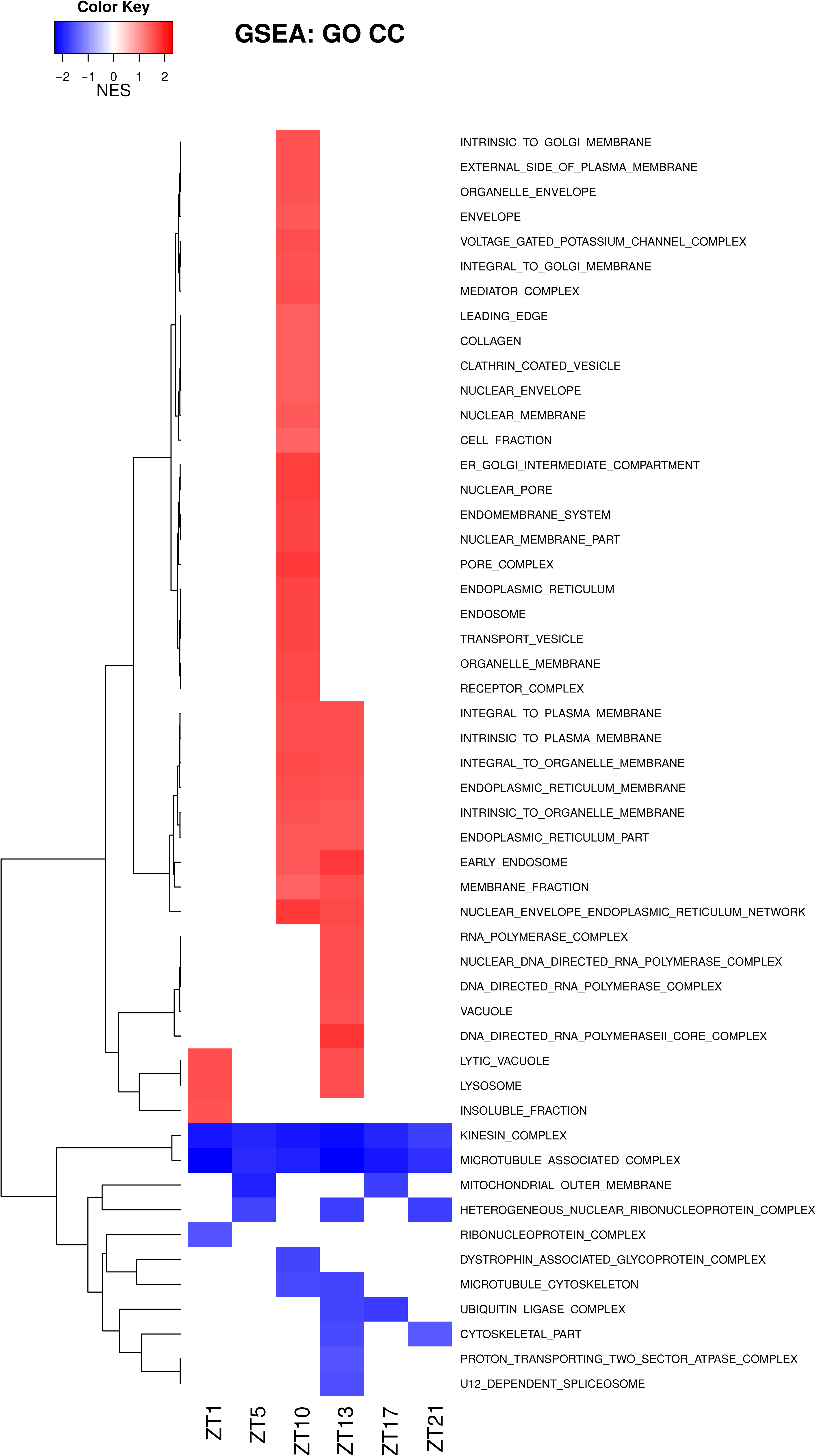

**figure s11.**
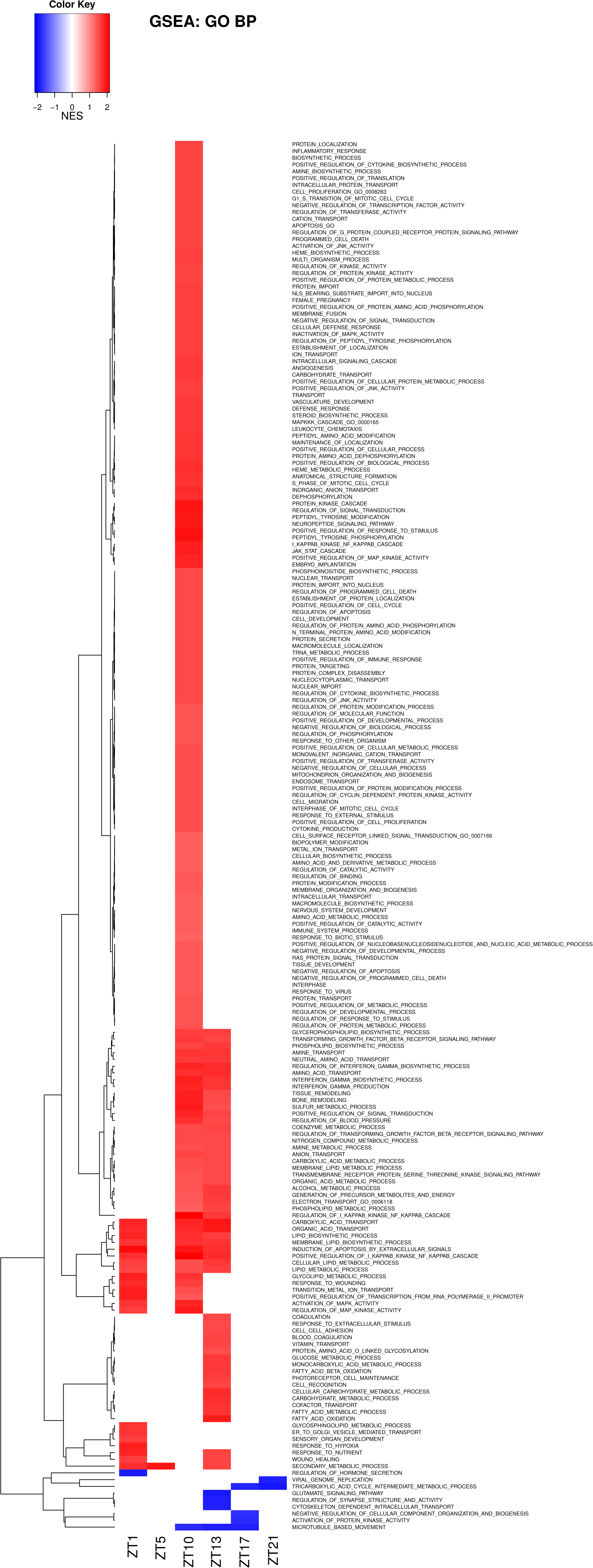

**figure s12.**
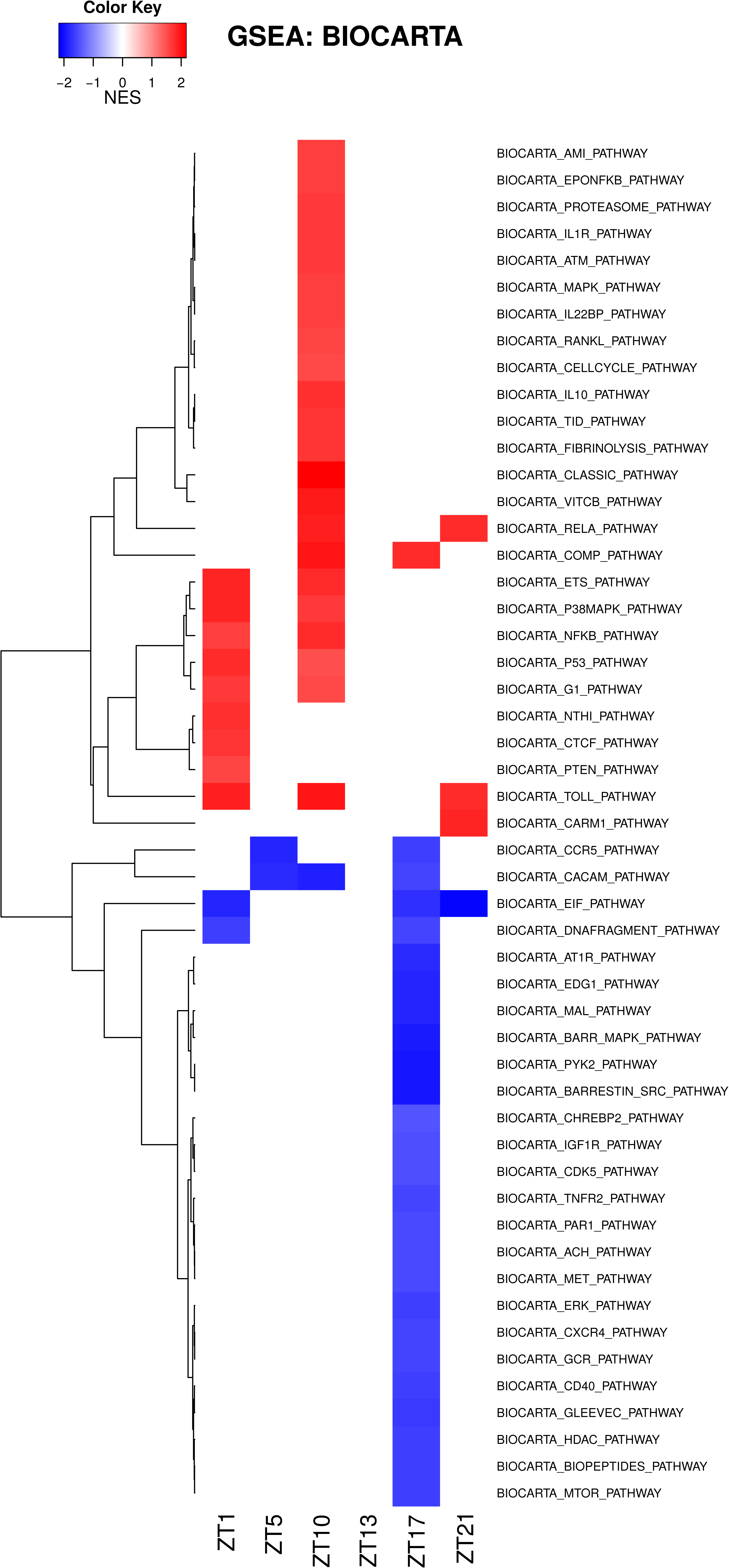

**figure s13.**
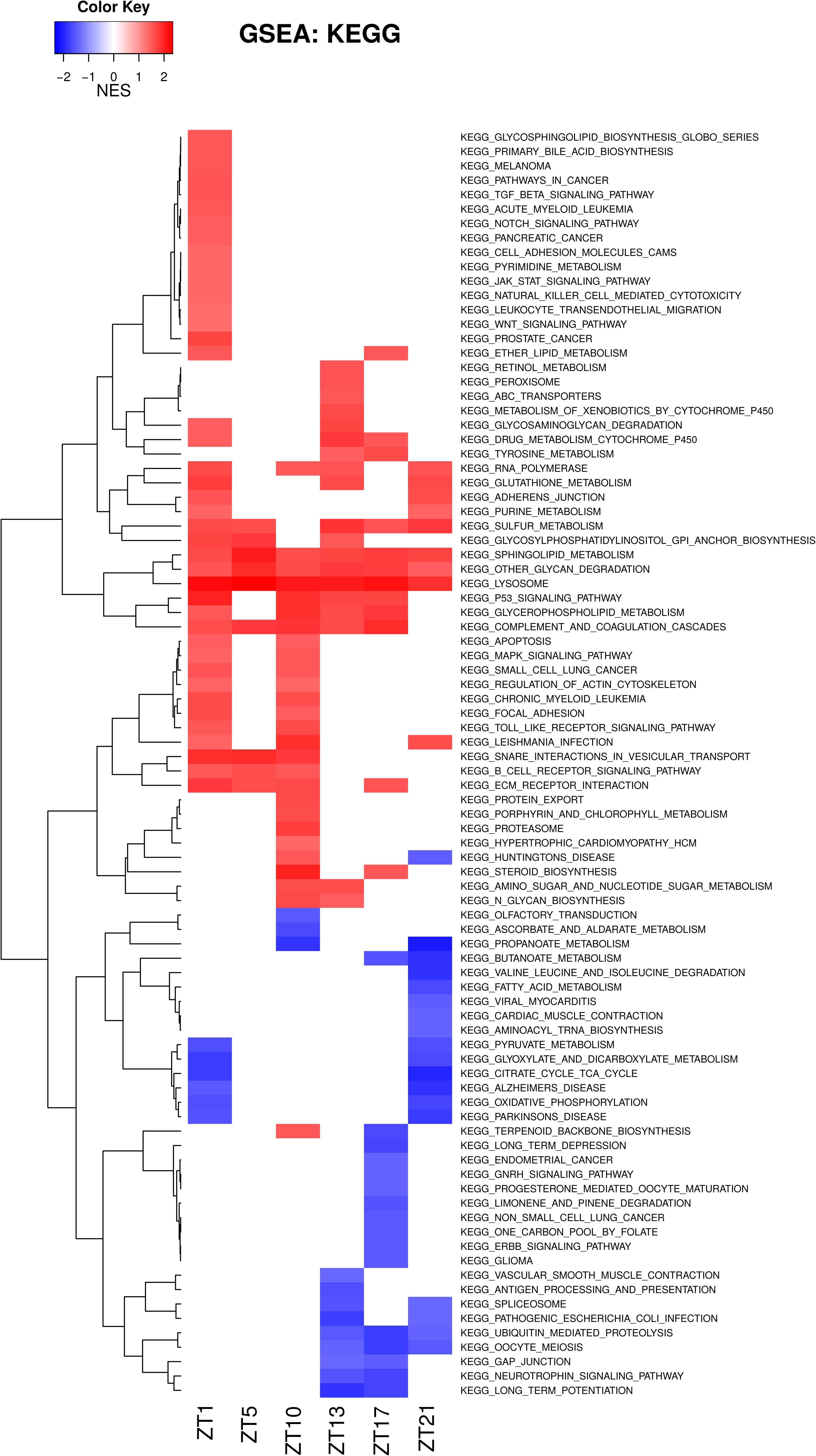

**figure s14.**
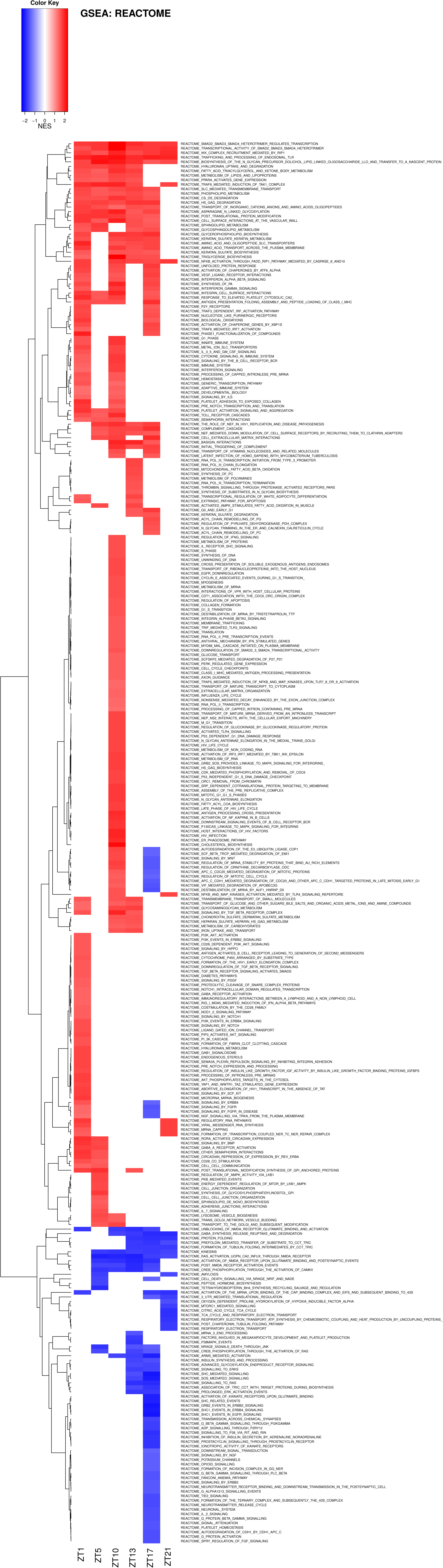

**figure S15.**
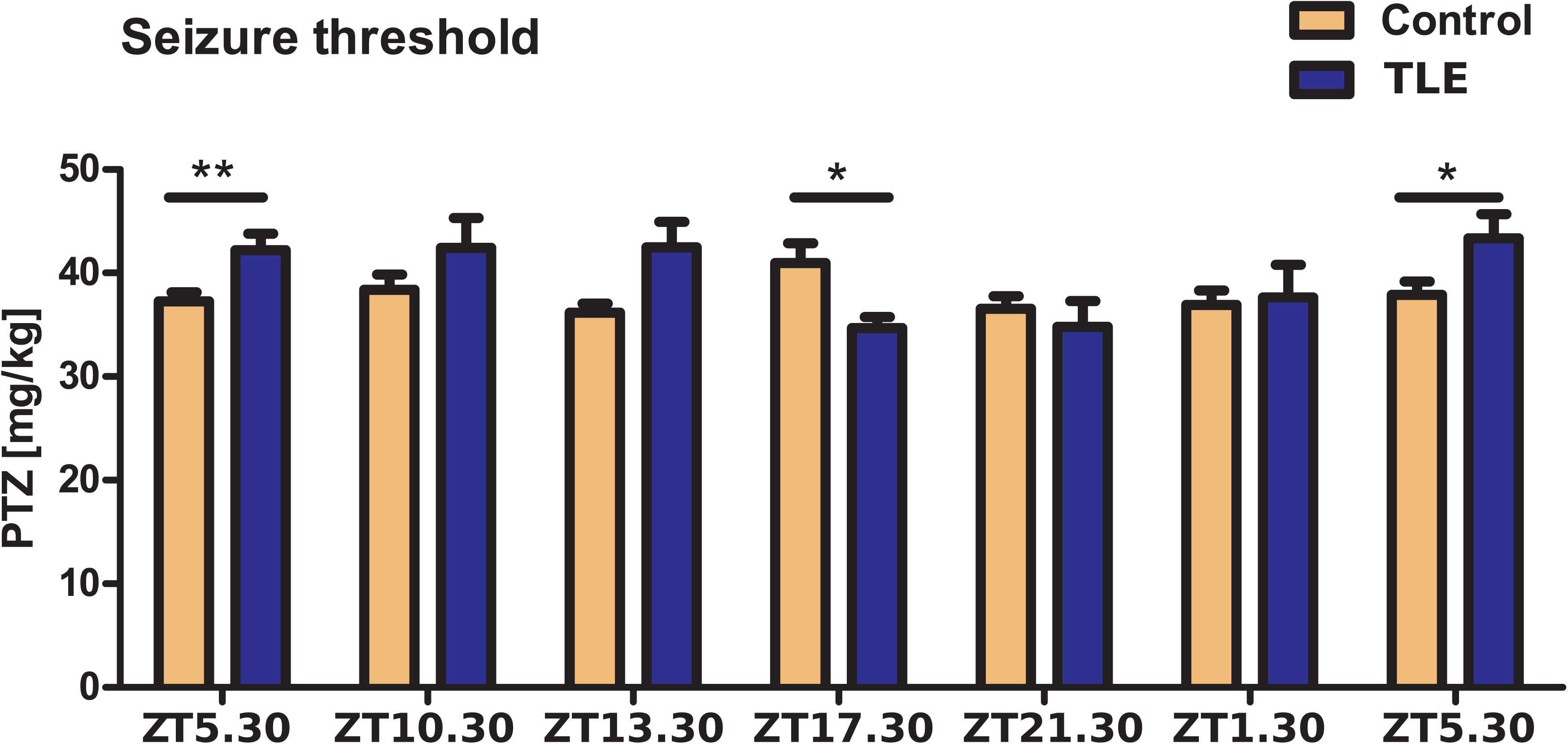
Thresholds of control mice did not differ significantly, whereas thresholds of TLE animals were significantly lower at ZT23.30 and ZT3.30 than during day-time represented by ZT11.30 (one way ANOVA P=0.0171, post hoc Bonferroni P<0.05). Both groups and time- points were compared using a two-way ANOVA, which revealed significant differences between the two groups at the time-points ZT11.30 and ZT23:30 (two-way ANOVA, followed by unpaired t-test *P<0.05;**P<0.01).

**Tables S1 to S4.** Results of transcriptome (S1) and proteome (S2) analyses for control and TLE (S3 and S4) animals, respectively.

**Tables S5 to S13.** Enriched terms reported from DAVID bioinformatics for oscillating genes found in control condition alone, epileptic condition alone, and oscillating in both conditions for 0.05 (S5 to S7), 0.01 (S8 to S10) and 0.001 (S11 to S13) p-values, respectively.

**Tables S14 to S25.** Raw data corresponding to figures S3 to S14 for Gene Ontology Molecular Function (GO_MF), Cellular Component (GO_CC), Biological Process (GO_BP), BIOCARTA, KEGG and REACTOME.

## References

1. Eckel-Mahan, K. & Sassone-Corsi, P. Metabolism and the circadian clock converge. Physiological reviews 93, 107–135 (2013).

2. Masri, S. & Sassone-Corsi, P. The circadian clock: a framework linking metabolism, epigenetics and neuronal function. Nature reviews. Neuroscience 14, 69–75 (2013).

3. Guilding, C. & Piggins, H.D. Challenging the omnipotence of the suprachiasmatic timekeeper: are circadian oscillators present throughout the mammalian brain? The European journal of neuroscience 25, 3195–3216 (2007).

4. Chun, L.E., Woodruff, E.R., Morton, S., Hinds, L.R. & Spencer, R.L. Variations in Phase and Amplitude of Rhythmic Clock Gene Expression across Prefrontal Cortex, Hippocampus, Amygdala, and Hypothalamic Paraventricular and Suprachiasmatic Nuclei of Male and Female Rats. Journal of biological rhythms 30, 417–436 (2015).

5. Harbour, V.L., Weigl, Y., Robinson, B. & Amir, S. Comprehensive mapping of regional expression of the clock protein PERIOD2 in rat forebrain across the 24-h day. PloS one 8, e76391 (2013).

6. Yang, S., Wang, K., Valladares, O., Hannenhalli, S. & Bucan, M. Genome-wide expression profiling and bioinformatics analysis of diurnally regulated genes in the mouse prefrontal cortex. Genome biology 8, R247 (2007).

7. Li, J.Z., et al. Circadian patterns of gene expression in the human brain and disruption in major depressive disorder. Proceedings of the National Academy of Sciences of the United States of America 110, 9950–9955 (2013).

8. Smarr, B.L., Jennings, K.J., Driscoll, J.R. & Kriegsfeld, L.J. A time to remember: the role of circadian clocks in learning and memory. Behav Neurosci 128, 283–303 (2014).

9. Gerstner, J.R., et al. Cycling behavior and memory formation. J Neurosci 29, 12824–12830 (2009).

10. Barnes, C.A., McNaughton, B.L., Goddard, G.V., Douglas, R.M. & Adamec, R. Circadian rhythm of synaptic excitability in rat and monkey central nervous system. Science 197, 91–92 (1977).

11. Harris, K.M. & Teyler, T.J. Age differences in a circadian influence on hippocampal LTP. Brain research 261, 69–73 (1983).

12. Chaudhury, D., Wang, L.M. & Colwell, C.S. Circadian regulation of hippocampal long-term potentiation. Journal of biological rhythms 20, 225–236 (2005).

13. Eckel-Mahan, K.L., et al. Circadian oscillation of hippocampal MAPK activity and cAmp: implications for memory persistence. Nat Neurosci 11, 1074–1082 (2008).

14. Zelinski, E.L., Deibel, S.H. & McDonald, R.J. The trouble with circadian clock dysfunction: multiple deleterious effects on the brain and body. Neuroscience and biobehavioral reviews 40, 80–101 (2014).

15. Orozco-Solis, R. & Sassone-Corsi, P. Epigenetic control and the circadian clock: linking metabolism to neuronal responses. Neuroscience 264, 76–87 (2014).

16. Karatsoreos, I.N. Links between Circadian Rhythms and Psychiatric Disease. Front Behav Neurosci 8, 162 (2014).

17. van Golde, E.G., Gutter, T. & de Weerd, A.W. Sleep disturbances in people with epilepsy; prevalence, impact and treatment. Sleep Med Rev 15, 357–368 (2011).

18. Eckel-Mahan, K.L., et al. Reprogramming of the circadian clock by nutritional challenge. Cell 155, 1464–1478 (2013).

19. Santos, E.A., et al. Diurnal Variation Has Effect on Differential Gene Expression Analysis in the Hippocampus of the Pilocarpine-Induced Model of Mesial Temporal Lobe Epilepsy. PloS one 10, e0141121 (2015).

20. Silver, R. & Kriegsfeld, L.J. Circadian rhythms have broad implications for understanding brain and behavior. The European journal of neuroscience 39, 1866–1880 (2014).

21. Zhang, R., Lahens, N.F., Ballance, H.I., Hughes, M.E. & Hogenesch, J.B. A circadian gene expression atlas in mammals: implications for biology and medicine. Proceedings of the National Academy of Sciences of the United States of America 111, 16219–16224 (2014).

22. Thompson, C.L., et al. Genomic anatomy of the hippocampus. Neuron 60, 1010–1021 (2008).

23. Marcelin, B., et al. Differential dorso-ventral distributions of Kv4.2 and HCN proteins confer distinct integrative properties to hippocampal CA1 pyramidal cell distal dendrites. The Journal of biological chemistry 287, 17656–17661 (2012).

24. Toyoda, I., Bower, M.R., Leyva, F. & Buckmaster, P.S. Early activation of ventral hippocampus and subiculum during spontaneous seizures in a rat model of temporal lobe epilepsy. The Journal of neuroscience: the official journal of the Society for Neuroscience 33, 11100–11115 (2013).

25. Magrane, M. & Consortium, U. UniProt Knowledgebase: a hub of integrated protein data. Database (Oxford) 2011, bar009 (2011).

26. Ivanov, A.I., Bernard, C. & Turner, D.A. Metabolic responses differentiate between interictal, ictal and persistent epileptiform activity in intact, immature hippocampus in vitro. Neurobiology of disease 75, 1–14 (2015).

27. Kann, O., et al. Metabolic dysfunction during neuronal activation in the ex vivo hippocampus from chronic epileptic rats and humans. Brain: a journal of neurology 128, 2396–2407 (2005).

28. Quigg, M., Straume, M., Menaker, M. & Bertram, E.H., 3rd. Temporal distribution of partial seizures: comparison of an animal model with human partial epilepsy. Annals of neurology 43, 748–755 (1998).

29. Jirsa, V.K., Stacey, W.C., Quilichini, P.P., Ivanov, A.I. & Bernard, C. On the nature of seizure dynamics. Brain: a journal of neurology 137, 2210–2230 (2014).

30. Law, V., et al. DrugBank 4.0: shedding new light on drug metabolism. Nucleic acids research 42, D1091–1097 (2014).

31. Rogawski, M.A., Loscher, W. & Rho, J.M. Mechanisms of Action of Antiseizure Drugs and the Ketogenic Diet. Cold Spring Harbor perspectives in medicine (2016).

32. Munn, R.G., Tyree, S.M., McNaughton, N. & Bilkey, D.K. The frequency of hippocampal theta rhythm is modulated on a circadian period and is entrained by food availability. Front Behav Neurosci 9, 61 (2015).

33. Munn, R.G. & Bilkey, D.K. The firing rate of hippocampal CA1 place cells is modulated with a circadian period. Hippocampus 22, 1325–1337 (2012).

34. Ly, J.Q., et al. Circadian regulation of human cortical excitability. Nature communications 7, 11828 (2016).

35. Wang, L.M., et al. Expression of the circadian clock gene Period2 in the hippocampus: possible implications for synaptic plasticity and learned behaviour. ASN neuro 1(2009).

36. Harbour, V.L., Weigl, Y., Robinson, B. & Amir, S. Phase differences in expression of circadian clock genes in the central nucleus of the amygdala, dentate gyrus, and suprachiasmatic nucleus in the rat. PloS one 9, e103309 (2014).

37. Prolo, L.M., Takahashi, J.S. & Herzog, E.D. Circadian rhythm generation and entrainment in astrocytes. The Journal of neuroscience: the official journal of the Society for Neuroscience 25, 404–408 (2005).

38. Kobow, K. & Blumcke, I. Epigenetic mechanisms in epilepsy. Prog Brain Res 213, 279–316 (2014).

39. McClelland, S., et al. Neuron-restrictive silencer factor-mediated hyperpolarization-activated cyclic nucleotide gated channelopathy in experimental temporal lobe epilepsy. Annals of neurology 70, 454–464 (2011).

40. Musiek, E.S. & Holtzman, D.M. Mechanisms linking circadian clocks, sleep, and neurodegeneration. Science 354, 1004-1008 (2016).

## References

1. Patel, V.R., Eckel-Mahan, K., Sassone-Corsi, P. & Baldi, P. CircadiOmics: integrating circadian genomics, transcriptomics, proteomics and metabolomics. Nature methods 9, 772–773 (2012).

42. Patel, V.R., et al. The pervasiveness and plasticity of circadian oscillations: the coupled circadian-oscillators framework. Bioinformatics 31, 3181–3188 (2015).

43. Bankstahl, M., Bankstahl, J.P. & Loscher, W. Pilocarpine-induced epilepsy in mice alters seizure thresholds and the efficacy of antiepileptic drugs in the 6-Hertz psychomotor seizure model. Epilepsy research 107, 205–216 (2013).

44. Becker, C., et al. Predicting and treating stress-Induced vulnerability to epilepsy and depression. Annals of neurology 78, 128–136 (2015).

45. Rattka, M., Brandt, C., Bankstahl, M., Broer, S. & Loscher, W. Enhanced susceptibility to the GABA antagonist pentylenetetrazole during the latent period following a pilocarpine-induced status epilepticus in rats. Neuropharmacology 60, 505–512 (2011).

46. Pollack, G.M. & Shen, D.D. A timed intravenous pentylenetetrazol infusion seizure model for quantitating the anticonvulsant effect of valproic acid in the rat. Journal of pharmacological methods 13, 135–146 (1985).

47. Loscher, W. Preclinical assessment of proconvulsant drug activity and its relevance for predicting adverse events in humans. European journal of pharmacology 610, 1–11 (2009).

48. Bot, A.M., Debski, K.J. & Lukasiuk, K. Alterations in miRNA levels in the dentate gyrus in epileptic rats. PloS one 8, e76051 (2013).

49. Carvalho, B.S. & Irizarry, R.A. A framework for oligonucleotide microarray preprocessing. Bioinformatics 26, 2363–2367 (2010).

50. Hughes, M.E., Hogenesch, J.B. & Kornacker, K. JTK_CYCLE: an efficient nonparametric algorithm for detecting rhythmic components in genome-scale data sets. Journal of biological rhythms 25, 372–380 (2010).

51. Kayala, M.A. & Baldi, P. Cyber-T web server: differential analysis of high-throughput data. Nucleic acids research 40, W553–559 (2012).

52. Huang da, W., Sherman, B.T. & Lempicki, R.A. Systematic and integrative analysis of large gene lists using DAVID bioinformatics resources. Nature protocols 4, 44–57 (2009).

53. Huang da, W., Sherman, B.T. & Lempicki, R.A. Bioinformatics enrichment tools: paths toward the comprehensive functional analysis of large gene lists. Nucleic acids research 37, 1–13 (2009).

54. Reimand, J., Arak, T. & Vilo, J. g:Profiler--a web server for functional interpretation of gene lists (2011 update). Nucleic acids research 39, W307–315 (2011).

55. Reimand, J., Kull, M., Peterson, H., Hansen, J. & Vilo, J. g:Profiler--a web-based toolset for functional profiling of gene lists from large-scale experiments. Nucleic acids research 35, W193–200 (2007).

56. Xie, X., Rigor, P. & Baldi, P. MotifMap: a human genome-wide map of candidate regulatory motif sites. Bioinformatics 25, 167–174 (2009).

57. Daily, K., Patel, V.R., Rigor, P., Xie, X. & Baldi, P. MotifMap: integrative genome-wide maps of regulatory motif sites for model species. BMC bioinformatics 12, 495 (2011).

58. Consortium, E.P. An integrated encyclopedia of DNA elements in the human genome. Nature 489, 57–74 (2012).

59. Kall, L., Storey, J.D., MacCoss, M.J. & Noble, W.S. Assigning significance to peptides identified by tandem mass spectrometry using decoy databases. J Proteome Res 7, 29–34 (2008).

60. Bernard, C., et al. Acquired dendritic channelopathy in temporal lobe epilepsy. Science 305, 532–535 (2004).

61. Mayevsky, A. & Rogatsky, G.G. Mitochondrial function in vivo evaluated by NADH fluorescence: from animal models to human studies. American journal of physiology. Cell physiology 292, C615–640 (2007).

62. Cahoy, J.D., et al. A transcriptome database for astrocytes, neurons, and oligodendrocytes: a new resource for understanding brain development and function. The Journal of neuroscience: the official journal of the Society for Neuroscience 28, 264–278 (2008).

